# Identification of disease treatment mechanisms through the multiscale interactome

**DOI:** 10.1101/2020.04.30.069690

**Authors:** Camilo Ruiz, Marinka Zitnik, Jure Leskovec

## Abstract

Most diseases disrupt multiple proteins, and drugs treat such diseases by restoring the functions of the disrupted proteins. How drugs restore these functions, however, is often unknown as a drug’s therapeutic effects are not limited only to the proteins that the drug directly targets. Here, we develop the multiscale interactome, a powerful approach to explain disease treatment. We integrate disease-perturbed proteins, drug targets, and biological functions into a multiscale interactome network, which contains 478,728 interactions between 1,661 drugs, 840 diseases, 17,660 human proteins, and 9,798 biological functions. We find that a drug’s effectiveness can often be attributed to targeting proteins that are distinct from disease-associated proteins but that affect the same biological functions. We develop a random walk-based method that captures how drug effects propagate through a hierarchy of biological functions and are coordinated by the protein-protein interaction network in which drugs act. On three key pharmacological tasks, we find that the multiscale interactome predicts what drugs will treat a given disease more effectively than prior approaches, identifies proteins and biological functions related to treatment, and predicts genes that interfere with treatment to alter drug efficacy and cause serious adverse reactions. Our results indicate that physical interactions between proteins alone are unable to explain the therapeutic effects of drugs as many drugs treat diseases by affecting the same biological functions disrupted by the disease rather than directly targeting disease proteins or their regulators. We provide a general framework for identifying proteins and biological functions relevant in treatment, even when drugs seem unrelated to the diseases they are recommended for.

---

Complex diseases, like cancer, disrupt dozens of proteins that interact in underlying biological networks [1–4]. Treating such diseases requires practical means to control the networks that underlie the disease [5–7]. By targeting even a single protein, a drug can affect hundreds of proteins in the underlying biological network. To achieve this effect, the drug relies on physical interactions between proteins. The drug binds a target protein, which physically interacts with dozens of other proteins, which in turn interact with dozens more, eventually reaching the proteins disrupted by the disease [8–10]. Networks capture such interactions and are a powerful paradigm to investigate the intricate effects of disease treatments and how these treatments translate into therapeutic benefits, revealing insights into drug efficacy [10–15], side effects [16], and effective combinatorial therapies for treating the most dreadful diseases, including cancers and infectious diseases [17–19].

However, existing systematic approaches assume that, for a drug to treat a disease, the proteins targeted by the drug need to be *close* to or even need to *coincide* with the disease-perturbed proteins [10–14] (Figure 1). As such, current approaches fail to capture biological functions, through which target proteins can restore the functions of disease-perturbed proteins and thus treat a disease [20–25] (Supplementary Fig. 3). Moreover, current systematic approaches are “black-boxes:” they predict treatment relationships but provide little biological insight into how treatment occurs. This suggests an opportunity for a systematic, explanatory approach. Indeed for particular drugs and diseases, custom networks have demonstrated that incorporating specific biological functions can help explain treatment [26–29].

**Figure 1:**
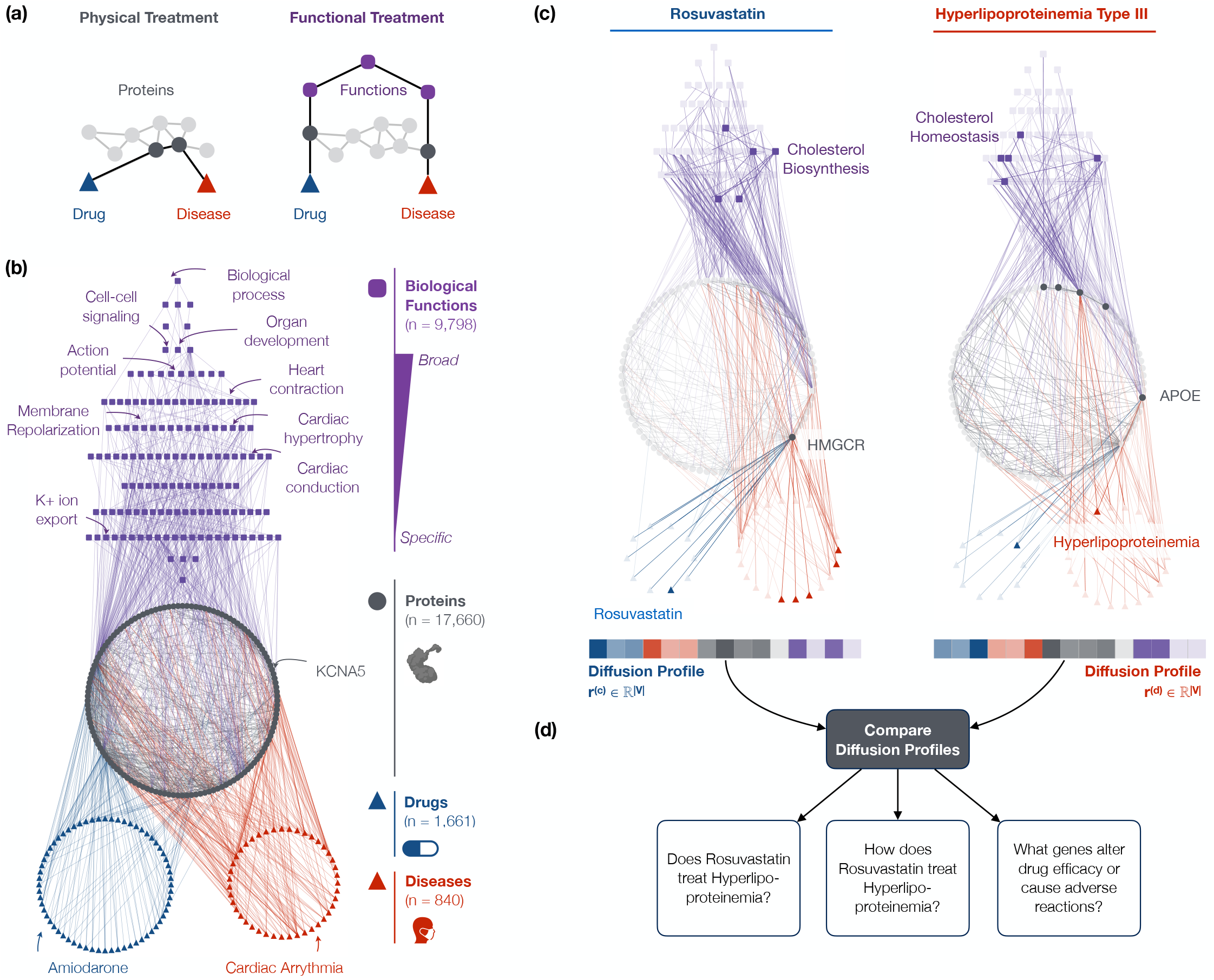
The multiscale interactome models drug treatment through both proteins and biological functions. **(a)** Existing systematic network approaches assume that drugs treat diseases by targeting proteins that are proximal to disease proteins in a network of physical interactions [10–14]. However, drugs can also treat diseases by targeting distant proteins that affect the same biological functions (Supplementary Fig. 3) [20–25]. **(b)** The multiscale interactome models drug-disease treatment by integrating both proteins and a hierarchy of biological functions (Supplementary Fig. 1). **(c)** The diffusion profile of a drug or disease captures its effect on every protein and biological function. The diffusion profile propagates the effect of the drug or disease via random walks which adaptively explore proteins and biological functions based on optimized edge weights. Ultimately, the visitation frequency of a node corresponds to the drug or disease’s propagated effect on that node (Methods). **(d)** By comparing the diffusion profiles of a drug and disease, we compare their effects on both proteins and biological functions. Thereby, we predict whether the drug treats the disease (Figure 2a-c), identify proteins and biological functions related to treatment (Figure 2d-h), and identify which genes alter drug efficacy or cause dangerous adverse reactions (Figure 3). For example, Hyperlipoproteinemia Type III’s diffusion profile reveals how defects in APOE affect cholesterol homeostasis, a hallmark of the excess blood cholesterol found in patients [49–53]. The diffusion profile of Rovustatin, a treatment for Hyperlipoproteinemia Type III, reveals how binding of HMG-CoA Reductase (HMGCR) reduces the production of excess cholesterol [54, 55]. By comparing these diffusion profiles, we thus predict that Rosuvastatin treats Hyperlipoproteinemia Type III, identify the HMGCR and APOE-driven cholesterol metabolic functions relevant to treatment, and predict that mutations in APOE and HMGCR may interfere with treatment and thus alter drug efficacy or cause dangerous adverse reactions.

Here we present the multiscale interactome, a powerful approach to explain disease treatment. We integrate disease-perturbed proteins, drug targets and biological functions in a multiscale interactome network. The multiscale interactome uses the physical interaction network between 17,660 human proteins, which we augment with 9,798 biological functions, in order to fully capture the fundamental biological principles of effective treatments across 1,661 drugs and 840 diseases.

To identify how a drug treats a disease, our approach uses biased random walks which model how drug effects spread through a hierarchy of biological functions and are coordinated by the protein-protein interaction network in which drugs act. In the multiscale interactome, drugs treat diseases by propagating their effects through a network of physical interactions between proteins and a hierarchy of biological functions. For each drug and disease, we learn a diffusion profile, which identifies the key proteins and biological functions involved in a given treatment. By comparing drug and disease diffusion profiles, the multiscale interactome provides an interpretable basis to identify the proteins and biological functions that explain successful treatments.

We demonstrate the power of the multiscale interactome on three key tasks in pharmacology. First, we find the multiscale interactome predicts which drugs can treat a given disease more accurately than existing methods that rely on physical interactions between proteins (i.e. a molecular-scale interactome). This finding indicates that our approach accurately captures the biological functions through which target proteins affect the functions of disease-perturbed proteins, even when drugs are distant to diseases they are recommended for. The multiscale interactome also improves prediction on entire drug classes, such as hormones, that rely on biological functions and thus cannot be accurately represented by approaches which only consider physical interactions between proteins. Second, we find that the multiscale interactome is a “white-box” method with the ability to identify proteins and biological functions relevant in treatment. Finally, we find that the multiscale interactome predicts what genes alter drug efficacy or cause serious adverse reactions for a given treatment and identifies biological functions that help explain how these genes interfere with treatment.

Our results indicate that the failure of existing approaches is not due to algorithmic limitations but is instead fundamental. We find that a drug can treat a disease by influencing the behaviors of proteins that are *distant* from the drug’s direct targets in the protein-protein interaction network. We find evidence that as long as those proteins affect the same biological functions disrupted by the disease proteins, the treatment can be successful. Thus, physical interactions between proteins alone are unable to explain the therapeutic effects of drugs, and functional information provides an important component for modeling treatment mechanisms. We provide a general framework for identifying proteins and biological functions relevant in treatment, even when drugs seem unrelated to the diseases they are recommended for.

## Results

### The multiscale interactome represents the effects of drugs and diseases on proteins and biological functions

The multiscale interactome models drug treatment by integrating both physical interactions between proteins and a multiscale hierarchy of biological functions. Crucially, many treatments depend on biological functions (Supplementary Fig. 3) [20–24]. Existing systematic network approaches, however, primarily model physical interactions between proteins [10–14], and thus cannot accurately model such treatments (Figure 1a, Supplementary Fig. 1).

Our multiscale interactome captures the fact that drugs and diseases exert their effects through both proteins and biological functions (Figure 1b). In particular, the multiscale interactome is a network in which 1,661 drugs interact with the human proteins they primarily target (8,568 edges) [30, 31] and 840 diseases interact with the human proteins they disrupt through genomic alterations, altered expression, or post-translational modification (25,212 edges) [32]. Subsequently, these protein-level effects propagate in two ways. First, 17,660 proteins physically interact with other proteins according to regulatory, metabolic, kinase-substrate, signaling, and binding relationships (387,626 edges) [33–39]. Second, these proteins alter 9,798 biological functions according to a rich hierarchy ranging from specific processes (i.e. embryonic heart tube elongation) to broad processes (i.e. heart development). Biological functions can describe processes involving molecules (i.e. DNA demethylation), cells (i.e. the mitotic cell cycle), tissues (i.e. muscle atrophy), organ systems (i.e. activation of the innate immune response), and the whole organism (i.e. anatomical structure development) (34,777 edges between proteins and biological functions, 22,545 edges between biological functions; Gene Ontology) [40, 41]. By modeling the effect of drugs and diseases on both proteins and biological functions, our multiscale interactome can model the range of drug treatments that rely on both [20–24].

Overall, our multiscale interactome provides a large, systematic dataset to study drug-disease treatments. Nearly 6,000 approved treatments (i.e., drug-disease pairs) spanning almost every category of human anatomy are compiled [31, 42, 43], exceeding the largest prior network-based study by 10X [13] (Anatomical Therapeutic Classification; Supplementary Fig. 4).

### Propagation of the effects of drugs and diseases through the multiscale interactome

To learn how the effects of drugs and diseases propagate through proteins and biological functions, we harnessed network diffusion profiles (Figure 1c). A network diffusion profile propagates the effects of a drug or disease across the multiscale interactome, revealing the most affected proteins and biological functions. The diffusion profile is computed by biased random walks that start at the drug or disease node. At every step, the walker can restart its walk or jump to an adjacent node based on optimized edge weights. The diffusion profile **r** ∈ ℝ^|*V*|^ measures how often each node in the multiscale interactome is visited, thus encoding the effect of the drug or disease on every protein and biological function.

Diffusion profiles contribute three methodological advances. First, diffusion profiles provide a general framework to adaptively integrate physical interactions between proteins and a hierarchy of biological functions. When continuing its walk, the random walker jumps between proteins and biological functions at different hierarchical levels based on optimized edge weights. These edge weights encode the relative importance of different types of nodes: *w*_drug_, *w*_disease_, *w*_protein_, *w*_biological function_, *w*_higher-level biological function_, *w*_lower-level biological function_. These weights are hyperparameters which we optimize when predicting the drugs that treat a given disease (Methods). For drug and disease treatments, these optimized edge weights encode the knowledge that proteins and biological functions at different hierarchical levels have different importance in the effects of drugs and diseases [20, 21]. By adaptively integrating both proteins and biological functions in a hierarchy, therefore, diffusion profiles model effects that rely on both.

Second, diffusion profiles provide a mathematical formalization of the principles governing how drug and disease effects propagate in a biological network. Drugs and diseases are known to generate their effects by disrupting or binding to proteins which recursively affect other proteins and biological functions. The effect propagates via two principles [8, 9]. First, proteins and biological functions closer to the drug or disease are affected more strongly. Similarly in diffusion profiles, proteins and biological functions closer to the drug or disease are visited more often since the random walker is more likely to visit them after a restart. Second, the net effect of the drug or disease on any given node depends on the net effect on each neighbor. Similarly in diffusion profiles, a random walker can arrive at a given node from any neighbor.

Finally, comparing diffusion profiles provides a rich, interpretable basis to predict pharmacological properties. Traditional random walk approaches predict properties by measuring the proximity of drug and disease nodes [9]. By contrast, we compare drug and disease diffusion profiles to compare their effects on proteins and biological functions, a richer comparison. Our approach is thus consistent with recent machine learning advances which harness diffusion profiles to represent nodes [44].

### The multiscale interactome accurately predicts which drugs treat a disease

By comparing the similarity of drug and disease diffusion profiles, the multiscale interactome predicts what drugs treat a given disease up to 40% more effectively than molecular-scale interactome approaches (AUROC 0.705 vs. 0.620, +13.7%; Average Precision 0.091 vs. 0.065, +40.0%; Recall@50 0.347 vs. 0.264, +31.4%) (Figure 2a, b, Methods). Note that drug-disease treatment relationships are never directly encoded into our network. Instead, the multiscale interactome learns to effectively predict drug-disease treatment relationships it has never previously seen.

**Figure 2:**
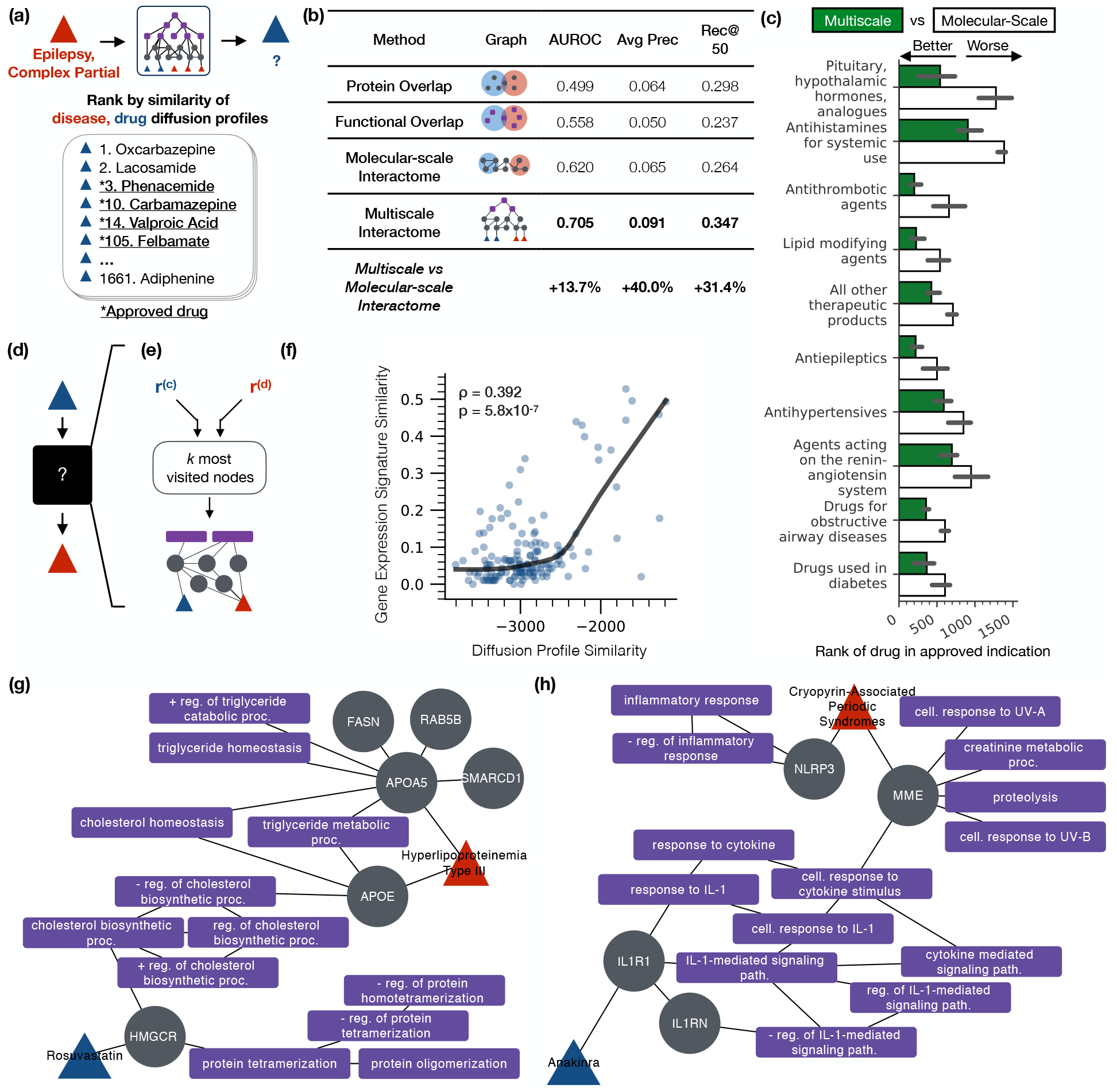
The multiscale interactome accurately predicts what drugs treat a disease and systematically identifies proteins and biological functions related to treatment. **(a)** To predict whether a drug treats a disease, we compare the drug and disease diffusion profiles according to a correlation distance. **(b)** By incorporating both proteins and biological functions, the multiscale interactome improves predictions of what drug will treat a given disease by up to 40% over molecular-scale interactome approaches [13]. Reported values are averaged across five-fold cross validation (Methods). **(c)** The multiscale interactome outperforms the molecular-scale interactome most greatly on drug classes that are known to harness biological functions which describe processes across the body (i.e., pituitary, hypothalamic hormones and analogues; median and 95% CI shown). **(d)** Existing interactome approaches are “black boxes”: they predict what drug treats a disease but do not explain how the drug treats the disease through specific biological functions [10–15]. **(e)** By contrast, the diffusion profiles of a drug and disease reveal the proteins and biological functions relevant to treatment. For each drug and disease pair, we induce a subgraph on the *k* most frequently visited nodes in the drug and disease diffusion profiles to explain treatment. **(f)** Drugs with more similar diffusion profiles have more similar gene expression signatures (Spearman *ρ* = 0.392, *p* = 5.8 × 10^−7^, *n* = 152), suggesting that drug diffusion profiles capture their biological effects. **(g)** The multiscale interactome explains treatments that molecular-scale interactome approaches cannot faith-fully represent. Rosuvastatin treats Hyperlipoprote Type III by binding to HMG CoA reductase (HMGCR) which drives a series of cholesterol biosynthetic functions affected by Hyperlipoproteinemia Type III [49–55]. **(h)** Anakinra treats Cryopyrin-Associated Periodic Syndromes by binding to IL1R1 which regulates immune-mediated inflammation through the Interleukin-1 beta signaling pathway [30, 57]. Inflammation is a hallmark of Cryopyrin-Associated Periodic Syndromes [56].

Moreover, the multiscale interactome accurately models classes of drugs that rely on biological functions and which molecular-scale interactome approaches thus cannot model effectively. Indeed, the top overall performing drug classes (i.e., sex hormones, modulators of the genital system; Supplementary Fig. 6) and the top drug classes for which the multiscale interactome out-performs the molecular-scale interactome (i.e., pituitary, hypothalamic hormones and analogues; Figure 2c, Supplementary Fig. 7) harness biological functions that describe processes across the body. For example, Vasopressin, a pituitary hormone, treats urinary disorders by binding receptors which trigger smooth muscle contraction in the gastrointestinal tract, free water reabsorption in the kidneys, and contraction in the vascular bed [30, 45, 46]. Treatment by Vasopressin, and by pituitary and hypothalamic hormones more broadly, relies on biological functions that describe processes across the body and that are modeled by the multiscale interactome.

### The multiscale interactome identifies proteins and biological functions relevant in complex treatments

Existing interactome approaches to systematically study treatment are “black boxes:” they predict what drug treats a disease but cannot explain how the drug treats the disease through specific proteins and biological functions [10–15] (Figure 2d). By contrast, drug and disease diffusion profiles identify proteins and biological functions relevant to treatment (Figure 2e, Supplementary Note 3). For a given drug and disease, we identify proteins and biological functions relevant to treatment by inducing a subgraph on the *k* most frequently visited nodes in the drug and disease diffusion profiles which correspond to the proteins and biological functions most affected by the drug and disease.

Gene expression signatures validate the biological relevance of diffusion profiles (Figure 2f). We find that drugs with more similar diffusion profiles have more similar gene expression signatures (Spearman *ρ* = 0.392, *p* = 5.8 × 10^−7^, *n* = 152) [47, 48], indicating that diffusion profiles reflect the effects of drugs on proteins and biological functions.

Furthermore, case studies validate the proteins and biological functions that diffusion profiles identify as relevant to treatment. Consider the treatment of Hyperlipoproteinemia Type III by Rosuvastatin (i.e., Crestor). In Hyperlipoproteinemia Type III, defects in apolipoprotein E (APOE) [49–51] and apolipoprotein A-V (APOA5) [52,53] lead to excess blood cholesterol, eventually leading to the onset of severe arteriosclerosis [50]. Rosuvastatin is known to treat Hyperlipoproteinemia Type III by inhibiting HMG-CoA reductase (HMGCR) and thereby diminishing cholesterol production [54, 55]. Crucially, diffusion profiles identify proteins and biological functions that recapitulate these key steps (Figure 2g). Notably, there is no direct path of proteins between Hyperlipoproteinemia and Rosuvastatin. Instead, treatment operates through biological functions (i.e., cholesterol biosynthesis and its regulation). Consistently, the multiscale interactome identifies Rosuvastatin as a treatment for Hyperlipoproteinemia far more effectively than a molecular-scale interactome approach, ranking Rosuvastatin in the top 4.33% of all drugs rather than the top 72.7%. The multiscale interactome explains treatments that rely on biological functions, a feat which molecular-scale interactome approaches cannot accomplish.

Similarly, consider the treatment of Cryopyrin-Associated Periodic Syndromes (CAPS) by Anakinra. In Cryopyrin-Associated Periodic Syndromes, mutations in NLRP3 and MME lead to immune-mediated inflammation through the Interleukin-1 beta signaling pathway [56]. Anakinra treats Cryopyrin-Associated Syndromes by binding IL1R1, a receptor which mediates regulation of the Interleukin-1 beta signaling pathway and thus prevents excessive inflammation [30,57]. Again, diffusion profiles identify proteins and biological functions that recapitulate these key steps (Figure 2h). Crucially, diffusion profiles identify the regulation of inflammation and immune system signaling, complex biological functions which are not modelled by molecular-scale interactome approaches. Again, the multiscale interactome identifies Anakinra as a treatment for CAPS far more effectively than a molecular-scale interactome approach, ranking Anakinra in the top 10.9% of all drugs rather than the top 71.8%.

### The multiscale interactome identifies genes that alter patient-specific drug efficacy and cause adverse reactions

A key goal of precision medicine is to understand how changes in genes alter patient-specific drug efficacy and cause adverse reactions [58] (Figure 3a). For particular treatments, detailed mechanistic models have been developed which can predict and explain drug resistance among genes already identified as relevant to treatment [26–29]. More systematically, however, current tools of precision medicine struggle to predict the genes that interfere with patient-specific treatment [59] and explain how such genes interfere with treatment [60].

**Figure 3:**
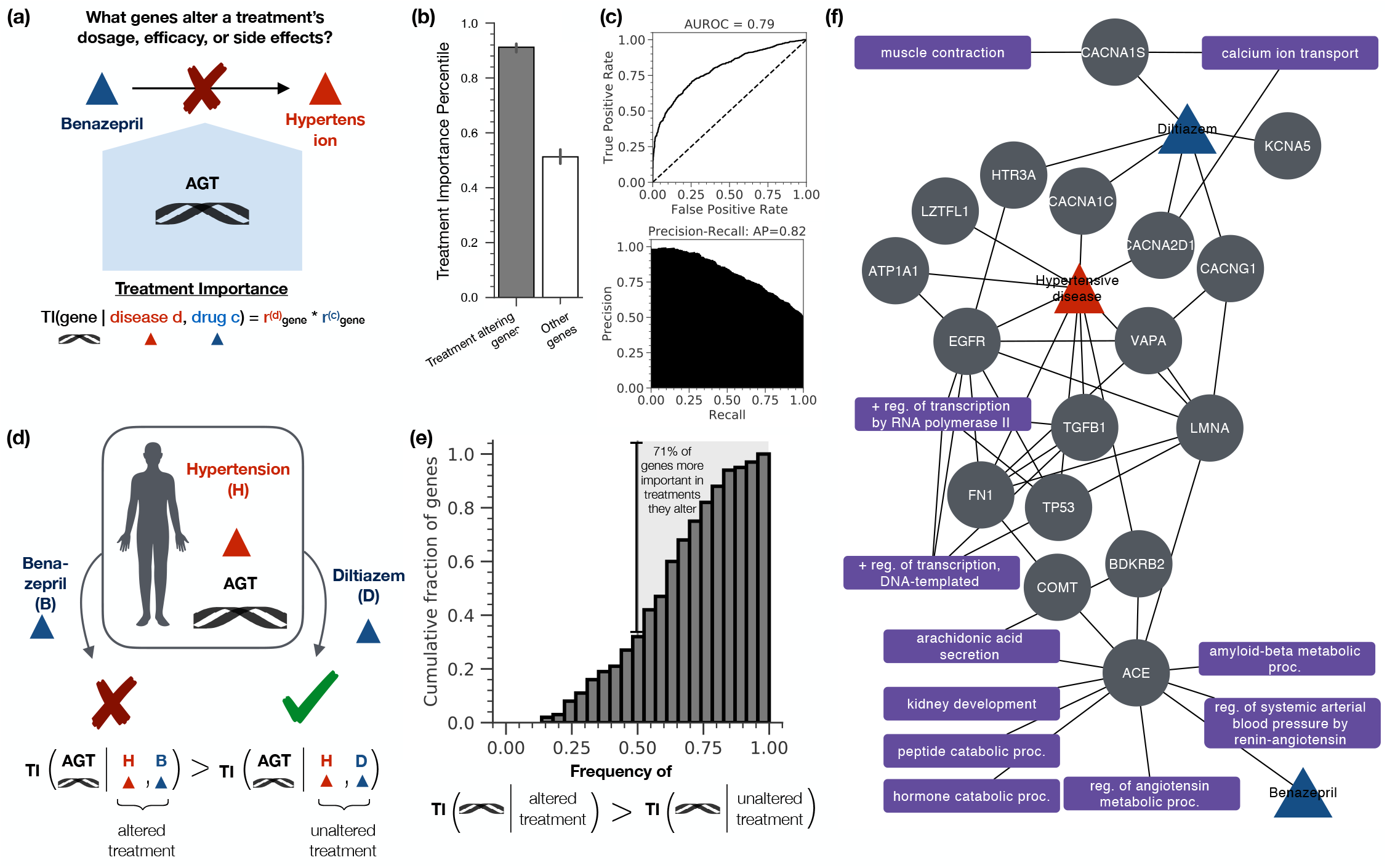
Diffusion profiles identify which genes alter drug efficacy and cause serious adverse reactions and identify biological functions that help explain the alteration in treatment. **(a)** Genes alter drug efficacy and cause serious adverse reactions in a range of treatments [61]. A pressing need exists to systematically identify genes that alter drug efficacy and cause serious adverse reactions for a given treatment and explain how these genes interfere with treatment [59]. **(b)** Genetic variants alter drug efficacy and cause serious adverse reactions by targeting genes of high network importance in treatment (median network importance of treatment altering genes = 0.912 vs. 0.513 *p* = 2.95 × 10^−107^; Mood’s median test; median and 95% CI shown). We define the network treatment importance of a gene according to its visitation frequency in the drug and disease diffusion profiles (Methods). **(c)** The treatment importance of a gene in the drug and disease diffusion profiles predicts whether that gene alters drug efficacy and causes serious adverse reactions for that particular treatment (AUROC = 0.79, Average Precision = 0.82). **(d)** Genes uniquely alter efficacy in one indicated drug but not another by primarily targeting the genes and biological functions used in treatment by the affected drug. In patients with Hypertensive Disease, a mutation in AGT alters the efficacy of Benazepril but not Diltiazem. Indeed, AGT exhibits a higher network importance in Benazepril treatment than in Diltiazem treatment, ranked as the 45th most important gene rather than the 418^th^ most important gene. **(e)** Overall, 71.0% of genes known to alter efficacy in one indicated drug but not another exhibit higher network importance in treatment by the affected drug. **(f)** Diffusion profiles can identify biological functions that may help explain alterations in treatment. Shown are the proteins and biological functions identified as relevant to the treatment of Hypertensive Disease by Benazepril and Diltiazem. AGT, which uniquely alters the efficacy of Benazepril, is a key regulator of the renin-angiotensin system, a biological function harnessed by Benazepril in treatment but not by Diltiazem [69–71].

We find that genetic variants that alter drug efficacy and cause serious adverse reactions occur in genes that are highly visited in the corresponding drug and disease diffusion profiles (Figure 3b). We define the treatment importance of a gene according to the visitation frequency of the corresponding protein in the drug and disease diffusion profiles (Methods). Genes that alter drug efficacy and cause adverse reactions exhibit substantially higher treatment importance scores than other genes (median network importance = 0.912 vs. 0.513; *p* = 2.95 × 10^−107^, Mood’s median test), indicating that these treatment altering genes occur at highly visited nodes. We thus provide evidence that the topological position of a gene influences its ability to alter drug efficacy or cause serious adverse reactions.

We find that the network importance of a gene in the drug and disease diffusion profiles predicts whether that gene alters drug efficacy and causes adverse reactions for that particular treatment (AUROC = 0.79, Average Precision = 0.82) (Figure 3c). Importantly, the knowledge that a gene alters a given treatment is never directly encoded into our network. Instead, diffusion profiles predict treatment altering relationships that the multiscale interactome has never previously seen. Our diffusion profiles thereby provide a systematic approach to identify genes with the potential to alter treatment. Our finding is complementary to high-resolution, temporal approaches such as discrete dynamic models which model drug resistance and adverse reactions by first curating genes and pathways deemed relevant to a particular treatment [26–29]. Diffusion profiles may help provide candidate genes and pathways for inclusion in these detailed approaches, including genes not previously expected to be relevant. New treatment altering genes, if validated experimentally and clinically, could ultimately affect patient stratification in clinical trials and personalized therapeutic selection [61].

Finally, we find that when a gene in a diseased patient alters the efficacy of one indicated drug but not another, that gene primarily targets the genes important to treatment for the resistant drug (Figure 3d, e). Overall, 71.0% of the genes known to alter the efficacy of one indicated drug but not another exhibit higher network importance in the altered treatments than in the unaltered treatment. We thus provide a network formalism explaining how changes to genes can alter efficacy and cause adverse reactions in only some drugs indicated to treat a disease.

Consider Benazepril and Diltiazem, two drugs indicated to treat Hypertensive Disease (Figure 3f). A mutation in the AGT gene alters the efficacy of Benazepril but not Diltiazem [62–64]. Indeed, our approach gives higher treatment importance to AGT in treatment by Benazepril than in treatment by Diltiazem, ranking AGT as the 45^th^ most important gene for Benazepril treatment but only the 418^th^ most important gene for Diltiazem treatment. Moreover, our approach explains why AGT alters the efficacy of Benazepril but not Diltiazem (Figure 3f). Diltiazem primarily operates at a molecular-scale, inhibiting various calcium receptors (CACNA1S, CACNA1C, CACNA2D1, CACNG1) which trigger relaxation of the smooth muscle lining blood vessels and thus lower blood pressure [30,65–67]. By contrast, Benazepril operates at a systems-scale: Benazepril binds to ACE which affects the renin-angiotensin system, a systems-level biological function that controls blood pressure through hormones [30,68,69]. Crucially, AGT or Angiotensinogen, is a key component of the renin-angiotensin system [69–71]. Therefore, AGT affects the key biological function used by Benazepril to treat Hypertensive Disease. By contrast, AGT plays no role in the calcium receptor driven pathways used by Diltiazem. Thus when a gene alters the efficacy of a drug, the multiscale interactome can identify biological functions that may help explain the alteration in treatment.

## Discussion

The multiscale interactome provides a general approach to systematically understand how drugs treat diseases. By integrating physical interactions and biological functions, the multiscale interactome improves prediction of what drugs will treat a disease by up to 40% over physical interactome approaches [10, 13]. Moreover, the multiscale interactome systematically identifies proteins and biological functions relevant to treatment. By contrast, existing systematic network approaches are “black-boxes” which make predictions without providing mechanistic insight. Finally, the multiscale interactome predicts what genes alter drug efficacy or cause severe adverse reactions for drug treatments and identifies biological functions that may explain how these genes interfere with treatment.

The multiscale interactome demonstrates that integrating biological functions into the interactome improves the systematic modeling of drug-disease treatment. Historically, systematic approaches to study treatment via the interactome have primarily focused on physical interactions between proteins [8–10, 13]. Here, we find that integrating biological functions into a physical interactome improves the systematic modeling of nearly 6,000 treatments. We find drugs and drug categories which depend on biological functions for treatment. More broadly, incorporating biological functions may improve systematic approaches that currently use physical interactions to study disease pathogenesis [72–75], disease comorbidities [6], and drug combinations [22–24]. Harnessing the multiscale interactome in these settings may thus help answer key pharmacological questions. Moreover, the multiscale interactome can be readily expanded to add additional node types relevant to the problem at hand (i.e. microRNAs to study cancer initiation and progression [76]). Our finding is consistent with systematic studies which demonstrate, in other contexts, that networks involving functional information can strengthen prediction of cellular growth [25, 77], identification of gene function [78–80], inference of drug targets [81], and general discovery of relationships between biological entities [82, 83].

Moreover, we find that diffusion profiles incorporating both proteins and biological functions provide predictive power and interpretability in modeling drug-disease treatments. Diffusion profiles predict what drugs treat a given disease and identify proteins and biological functions relevant to treatment. In other pharmacological contexts, diffusion profiles incorporating proteins and biological functions may thus improve systematic approaches which currently employ proximity or other non-interpretable methods [6, 16, 17, 33]. In studying the efficacy of drug combinations [17], diffusion profiles may identify synergistic effects on key biological functions. In studying the adverse reactions of drug combinations [16], diffusion profiles may identify biological functions which help explain polypharmacy side effects. In disease comorbidities [6, 33], diffusion profiles may predict new comorbidities and identify biological functions which help explain the development of the comorbidity.

Finally, our study shows that both physical interactions and biological functions can propagate the effects of drugs and diseases. We find that many drugs neither directly target the proteins associated with the disease they treat nor target proximal proteins. Instead, these drugs affect the same biological functions disrupted by the disease. This view expands upon the current view of indirect effects embraced in other biological phenomena. In the omnigenic model of complex disease [84, 85], for example, hundreds of genetic variants affect a complex phenotype through indirect effects that propagate through a regulatory network of physical interactions. Our results suggest that the multiscale interactome, incorporating both physical interactions and biological functions, may help propagate indirect effects in complex disease. Altogether, the multiscale interactome provides a general computational paradigm for network medicine.

## Data availability

All data used in the paper, including the multiscale interactome, approved drug-disease treatments, drug and disease classifications, gene expression signatures, and pharmacogenomic relationships is available at github.com/snap-stanford/multiscale-interactome.

## Code availability

Python implementation of our methodology is available at github.com/snap-stanford/multiscale-interactome. The code is written in Python. Please read the README for information on downloading and running the code.

## Author contributions

C.R., M.Z., and J.L. designed research; C.R., M.Z., and J.L. performed research; C.R., M.Z., and J.L. analyzed data; and C.R., M.Z., and J.L. wrote the paper.

## Corresponding author

Correspondence should be addressed to J.L.

## Methods

### The multiscale interactome

The multiscale interactome captures how drugs use both a network of physical interactions and a rich hierarchy of biological functions to treat diseases. In the multiscale interactome, 1,661 drugs connect to the proteins they target (8,568 edges) [30, 31]. 840 diseases connect to the proteins they disrupt through genomic alterations, altered expression, or post-translational modification (25,212 edges) [32]. 17,660 proteins connect to other proteins based on physical interactions such as regulatory, metabolic, kinase-substrate, signaling, or binding relationships (387,626 edges) [33–39]. Proteins connect to the 9,798 biological functions they affect (22,545 edges) [40, 41]. Finally, biological functions connect to each other in a rich hierarchy ranging from specific processes (i.e. embryonic heart tube elongation) to broad processes (i.e. heart development) (22,545 edges) [40, 41]. Biological functions can describe processes involving molecules (i.e. DNA demethylation), cells (i.e. the mitotic cell cycle), tissues (i.e. muscle atrophy), organ systems (i.e. activation of the innate immune response), and the whole organism (i.e. anatomical structure development).

We visualize a representative subset of the multiscale interactome using Cytoscape [86] (Figure 1b).

### Drug–protein interactions

We map drugs to their protein targets using DrugBank [30] and the Drug Repurposing Hub [31]. For DrugBank, we map the Uniprot Protein IDs to Entrez IDs using HUGO [87]. For the Drug Repurposing Hub, we map drugs to their DrugBank IDs using the drug names and DrugBank’s “drugbank_approved_target_uniprot_links.csv” file. We map protein targets to Entrez IDs using HUGO [87]. We filter drug-target relationships to only include proteins that are represented in the network of physical interactions between proteins (see Methods: Protein–protein interactions). All drug-target interactions are provided in Supplementary Data 1.

### Disease–protein interactions

We map diseases to genes they affect through genomic alterations, altered expression, or post-translational modification by using DisGeNet [32]. To ensure highquality disease-gene associations, we only consider the “curated” set of disease-gene associations provided by DisGeNet which draws from expert-curated repositories: UniProt, the Comparative Toxicogenomics Database, Orphanet, the Clinical Genome Resource (ClinGen), Genomics England PanelApp, the Cancer Genome Interpreter (CGI), and the Psychiatric Disorders Gene Association Network (PsyGeNET). We exclude all disease-gene associations that are inferred, based on orthology relationships from animal models, or based on computational-mining of the literature. Ultimately, diseases are associated with genes they affect via genomic alteration, alteration of expression, or post-translational modification according to the DisGeNet relationship ontology. To avoid circularity in the analysis, we remove disease-gene associations marked as therapeutic. Finally, we filter disease-gene relationships to only consider genes whose protein products were present in the network of physical interactions between proteins (see Methods: Protein–protein interactions). All disease-protein interactions are provided in Supplementary Data 2.

### Protein–protein interactions

We generate a network of 387,626 physical interactions between 17,660 proteins by compiling seven major databases. Across all databases, we only consider human proteins and their interactions; only allow protein-protein interactions with direct experimental evidence; and only allow *physical* interactions between proteins, filtering out genetic and indirect interactions between proteins such as those identified via synthetic lethality experiments. All protein-protein interactions are provided in Supplementary Data 3.

1. *The Biological General Repository for Interaction Datasets [34]* (BioGRID; 309,187 interactions between 16,352 proteins). BioGRID manually curates both physical and genetic interactions between proteins from 71,713 high- and low-throughput publications. We map BioGRID proteins to Entrez IDs by using HUGO [87]. We only include protein-protein interactions from BioGRID that result from experiments indicating a *physical* interaction between the proteins, as described by BioGRID [34], and ignore protein-protein interactions indicating a *genetic* interaction between the proteins. We use the “BIOGRID-ORGANISM-Homo sapiens-3.5.178.tab” file.
2. *The Database of Interacting Proteins [36]* (DIP; 4,235 interactions between 2,751 proteins). DIP only considers physical protein-protein interactions with experimental evidence and curates these from the literature. We map the UniProt ID of each protein to its Entrez ID by using HUGO [87]. We allow all experimental methods from DIP since they all capture physical interactions [36].
3. *The Human Reference Protein Interactome Mapping Project*. We integrate four protein-protein interaction networks from the Human Reference Protein Interactome Mapping Project that were generated through high-throughput yeast two hybrid assays (HI-I-05 [39]: 2,611 interactions between 1,522 proteins; HI-II-14 [35] 13,426 interactions between 4,228 proteins; Venkatesan-09 [37]: 233 interactions between 229 proteins; Yu-11 [38] 1,126 interactions between 1,126 proteins). Since protein-protein interactions in all four networks result from a yeast two-hybrid system, all protein-protein interactions are physical and experimentally verified. We thus include all protein-protein interactions across these networks. Proteins are already provided with their Entrez ID so no mapping is required.
4. *Menche-2015 [33]* (138,425 interactions between 13,393 proteins). Finally, we integrate the physical protein-protein interaction network compiled by Menche et al. [33]. Menche et al. compiles different types of physical protein-protein interactions from a range of sources. In all cases, protein-protein interactions result from direct experimental evidence. Menche et al. compiles regulatory interactions from the TRANSFAC database; binary interactions from a series of high-throughput yeast-two-hybrid datasets as well as the IntAct and MINT databases; literature curated interactions from IntAct, MINT, BioGRID, and HPRD; metabolic-enzyme coupled interactions from KEGG and BIGG; protein complex interactions from CORUM; kinase-substrate interactions from PhosphositePlus; and signaling interactions from [88]. All proteins are provided in Entrez format and thus do not require further mapping.

### Protein – biological function interactions

We map proteins to the biological functions they affect by using the human version of the Gene Ontology [40, 41] (7,993 proteins; 6,387 biological functions; 34,777 edges). We only allow experimentally verified associations between genes and biological functions according to the following IDs: EXP – inferred from experiment, IDA inferred from direct assay, IMP – inferred from mutant phenotype, IGI – inferred from genetic interaction, HTP – high throughput experiment, HDA – high throughput direct assay, HMP – high throughput mutant phenotype, and HGI – high throughput genetic interaction. We exclude any protein–biological function relationships that are inferred from physical interactions to avoid redundancy with the physical network of interacting proteins. We also exclude protein–biological function relationships inferred from gene expression patterns since the Gene Ontology states that such interactions are challenging to map to specific proteins [40, 41]. To prevent circularity, we further ignore all associations based on phylogenetically inferred annotations or various computational analyses (sequence or structural similarity, sequence orthology, sequence alignment, sequence modeling, genomic context, reviewed computational analysis). Finally, we ignore associations based on author statements, curator inference, electronic annotations (i.e. automated annotations), and those for which no biological data was available. Some biological functions in the Gene Ontology have multiple synonymous IDs. For each biological function, we use the “master IDs” provided by GOATOOLS [89]. All protein – biological function interactions are provided in Supplementary Data 4.

### Biological function – biological function interactions

We construct a hierarchy of biological functions by using the Gene Ontology’s Biological Processes [40, 41]. The Gene Ontology represents a curated hierarchy of biological functions, where highly specific biological functions are children of more general biological functions according to numerous relationship types. For example, “negative regulation of response to interferon-gamma” 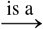 “negative regulation of innate immune response” 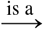 “negative regulation of immune response” 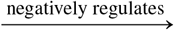 “immune response.” We allow relationships between biological functions of the following types: regulates, positively regulates, negatively regulates, part of, and is a. In order to allow the model to focus on the biological functions most relevant to treatment, we only consider biological functions which are associated with at least one drug target or one disease protein, either directly or implicitly through their children. All biological function – biological function interactions are provided in Supplementary Data 5.

### Constructing dataset of approved drug-disease treatments

We construct a dataset of 5,926 unique, approved drug-disease pairs, exceeding the largest prior network-based study by 10X [13]. We source approved drug-disease pairs from the Drug Repurposing Database [42] (*n*_*pairs*_ = 2, 538; *n*_*drugs*_ = 996, *n_diseases_* = 463), the Drug Repurposing Hub [31] (*n_pairs_* = 1, 449; *n_drugs_* = 908, *n_diseases_* = 265), and the Drug Indication Database [43] (*n_pairs_* = 3, 304; *n_drugs_* = 1, 147, *n_diseases_* = 615). In all cases, we filter drug-disease pairs to ensure that only FDA-approved treatment relationships are included.

We extract approved drug-disease pairs from each database as follows. In all cases, drugs are mapped to DrugBank IDs [30] and diseases are mapped to unique identifiers from the National Library of Medicine [90] (NLM UMLS CUIDs: NLM Unified Medical Language System Controlled Unique Identifier):

1. *The Drug Repurposing Database* is a gold-standard database of drug-disease pairs extracted from drug labels and the American Association of Clinical Trials Database [42]. Drugs and diseases in the Drug Repurposing Database are provided with DrugBank IDs and NLM UMLS CUIDs so no additional mapping is required. We extract only the drug and disease pairs designated as “Approved” treatment relationships.
2. *The Broad Institute’s Drug Repurposing Hub* is a hand-curated collection of drug-disease pairs compiled from drug labels, DrugBank, the NCATS NCGC Pharmaceutical Collection (NPC), Thomson Reuters Integrity, Thomson Reuters Cortellis, Citeline Pharmaprojects, the FDA Orange Book, ClinicalTrials.gov, and PubMed [31]. We map drugs to DrugBank IDs by comparing their provided names and PubChem IDs to DrugBank’s external links mapping [30]. We map diseases to UMLS CUIDs by using the UMLS Metathesaurus’s REST API [90]. Finally, we only include drug-disease pairs with a “Launched” clinical phase attribute, indicating FDA approval.
3. *The Drug Indication Database* provides drug-indications relationships from DailyMed, DrugBank, the Pharmacological Actions sections of the Medical Subject Headings, the National Drug File Reference Terminology, the Physicians’ Desk Reference, the Chemical Entities of Biological Interest (ChEBI), the Comparative Toxicogenomics Database, the Therapeutic Claims section of the USP Dictionary of United States Adopted Names and International Drug Names, and the World Health Organization Anatomic-Therapeutic-Chemical classification) [43]. The Drug Indication Database captures both diseases and non-disease medical conditions (i.e. pregnancy) for which a drug is used. Additionally, the Drug Indication Database captures both treatment relationships between drugs and indications as well as prevention, management, and diagnostic relationships. We filter the Drug Indication Database to only include *approved* treatment relationships between drugs and *diseases*. We map drugs to DrugBank IDs by using the provided CAS and ChEBI IDs as well as DrugBank’s external links mapping [30]. Indications are already provided with UMLS CUIDs. We filter indications to only include diseases in two ways. First, we only consider indications with a UMLS semantic type of “B2.2.1.2.1 Disease or Syndrome”, “B2.2.1.2 Pathologic Function”, or “B2.2.1.2.1.2 Neoplastic Process.” Second, we only consider indications present in DisGeNet, a database mapping diseases to their associated genes [32]. To ensure that drug-disease relationships specifically represent treatment relationships, we filter drug-disease pairs based on the “indication subtype.” We remove drug-indication pairs where the indication subtype described is not treatment (i.e. preventative/prophylaxis, diagnosis, adjunct, palliative, reduction, causes/inducing/associated, and mechanism). We additionally remove all drug indication pairs from the Comparative Toxicogenomics Database (CTD). The goal of CTD is to provide broad chemical-disease associations published in the literature [91]. Concurrently, CTD does not subset these chemical-disease associations into drug-disease relationships that represent FDA-approved treatments. Finally, we remove overly broad diseases from the Drug Indication Database. We remove disease categories (i.e. diseases with “Diseases” in their name such as “Cardiovascular Diseases” and “Metabolic Diseases). We also remove diseases with more than 130 approved drugs (i.e. Disorder of Eye – 290 approved drugs).

After compiling approved drug-disease treatment pairs, we remove treatments for which drugs rely on binding to non-human proteins (i.e. viral or bacterial proteins) to induce their effect. The multiscale interactome only models human proteins and biological functions. The multiscale interactome is thus not designed to model treatments which rely on binding to viral or bacterial proteins. To remove such treatments, we map all disease UMLS CUIDs to their corresponding Disease Ontology ID [92]. We then remove diseases corresponding to the “disease by infectious agent category” of the Disease Ontology. The Disease Ontology does not map many UMLS CUIDs to corresponding Disease Ontology IDs. We thus manually curate the final list of diseases to remove additional infectious diseases: malaria, bacterial septicemia, fungal infection, coccidiosis, gonorrhea, gastrointestinal roundworms, shingles, lice, gastrointestinal parasites, tapeworm, syphilis, genital herpes, lungworms, fungicide, fungal keratosis, yeast infection, laryngitis, enterocolitis, protozoan infection, African trypanosomiasis, sepsis, Chagas disease, mites, bacterial vaginosis, scabies, pinworm, equine protozoal myeloencephalitis (EPM), microsporidiosis, and ringworm.

Finally, we filter approved drug-disease treatment pairs to only include drugs with at least one known target in DrugBank [30] or the Drug Repurposing Hub [31] and diseases with at least one associated gene in the curated version of DisGeNet [32] as these are the only drugs and diseases that the multiscale interactome represents (see Methods: Drug–protein interactions, Disease–protein interactions).

Ultimately, we achieve a dataset of 5,926 approved drug-disease pairs, exceeding the largest prior network-based study by 10X [13]. All approved drug-disease pairs are provided in Supplementary Data 6.

### Learning drug and disease diffusion profiles

We propagate the effects of each drug and disease across the multiscale interactome by using network diffusion profiles. A drug or disease diffusion profile learns the proteins and biological functions most affected by each drug or disease. Each drug or disease diffusion profile is computed through biased random walks that start at the drug or disease node. At every step, the random walker can restart its walk or jump to an adjacent node based on optimized edge weights. After many walks, the diffusion profile measures how often every node was visited, thus representing the effect of the drug or disease on that node.

By using optimized edge weights, diffusion profiles learn to adaptively integrate proteins and biological functions. Diffusion profiles rely on a set of scalar weights which encode the relative importance of different types of nodes: *W* = {*w*_drug_, *w*_disease_, *w*_protein_, *w*_biological function_, *w*_higher-level biological function_, *w*_lower-level biological function_}. These weights are hyperparameters which we optimize when predicting the drugs that treat a given disease (see Methods: Model selection and optimization of scalar weights). When a random walker continues its walk, it picks the next node to jump to based on the relative values of these weights. For example, if a random walker is at a protein and has both protein and biological function neighbors, it is 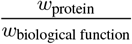 times more likely to jump to the protein neighbors than the biological function neighbors. Notice that proteins connect to drugs, diseases, proteins, and biological functions, making {*w*_drug_, *w*_disease_, *w*_protein_, *w*_biological function_} the relevant weights for a random walker currently at a protein. By contrast, biological functions connect to proteins, higher-level biological functions, and lower-level biological functions, making {*w*_protein_, *w*_higher-level biological function_, *w*_lower-level biological function_} the relevant weights for a random walker at a biological function. By providing separate weights for higher- and lower-level biological functions, the random walker learns to explore different levels of the hierarchy of biological functions and integrate them appropriately.

Diffusion profiles represent a general methodology to propagate signals through a heterogeneous biological network. By carefully defining edge weights and the nodes that the random walker restarts to, diffusion profiles can be used in a wide range of biological tasks. Here, we define edge weights for drug, disease, protein, and biological function node types, yet more or fewer weights can be used based on the problem of interest. Similarly, here, the random walker jumps to the initial drug or disease node after a restart, but in reality, it can restart to any node or any set of nodes. The edge weights and restart nodes thus make diffusion profiles a flexible approach to propagate signals across a heterogeneous biological network, with applicability to a wide range of problems in systems biology and pharmacology.

### Computing drug and disease diffusion profiles through power iteration

Mathematically, we compute diffusion profiles through a matrix formulation with power iteration [93–95]. The diffusion profile computation takes as input:

1. *G* = (*V, E*) the unweighted, undirected multiscale interactome with *V* nodes and *E* edges.
2. *W* = {*w*_drug_, *w*_disease_, *w*_protein_, *w*_biological function_, *w*_higher-level biological function_, *w*_lower-level biological function_} the set of scalar weights which encode the relative likelihood of the walker jumping from one node type to another when continuing its walk.
3. *α* which represents the probability of the walker continuing its walk at a given step rather than restarting.
4. **s** ∈ ℝ ^|*V*|^ a restart vector which sets the probability the walker will jump to each node after a restart; here, **s**is a one-hot vector encoding the drug or disease of interest.
5. *∊* the tolerance allowed for convergence of the power iteration computation.

The diffusion profile computation outputs **r** ∈ ℝ ^|*V*|^, a drug- or disease-diffusion profile which measures the frequency with which the random walker visits each node. Note that ∑_*i*_ **r**_*i*_ = 1.

Before computing the diffusion profile of a drug or disease of interest, we preprocess the multiscale interactome in order to only allow biologically meaningful walks. Diffusion profiles are designed to capture how a drug or disease of interest propagates its effect by recursively affecting proteins and biological functions. Notice that drugs and diseases do not propagate their effect by using other drugs and diseases as intermediates. Therefore, we disallow paths that have drugs and diseases as intermediate nodes. To accomplish this mathematically, we convert *G* = (*V, E*) to a directed graph *G′* where all previously undirected edges are replaced by edges in both directions (i.e. edges now include drug↔protein, disease↔protein, protein↔protein, protein↔biological function, and lower-level biological function↔higher-level biological function). We then make the drug or disease of interest a source node (i.e. no in-edges) and all other drugs and diseases sink nodes (i.e. no out-edges). In *G′*, a random walker starts at the drug or disease of interest and recursively walks to proteins and biological functions. If the walker reaches any other drug or disease node, it must restart its walk.

Next, we encode *G′* and the set of scalar weights *W* into a biased transition matrix **M** ∈ ℝ ^|*V*|×|*V*|^. Each entry **M**_*ij*_ denotes the probability *p*_*i*→*j*_ a random walker jumps from node *i* to node *j* when continuing its walk. Consider a random walker at node *i* jumping to neighbor *j* of type *t*. Let *T* be the set of all node types adjacent to node *i*. We compute *p*_*i*→*j*_ in two steps.

1. First, we compute the probability of the random walker jumping to a node of type *t* rather than a node of a different type. *w*_*t*_ is the weight of node type *t* as specified in *W*:

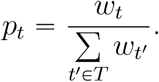
2. Second, we compute the probability that the random walker jumps to node *j* rather than to another adjacent node of type *t*. Let *n_t_* be the number of adjacent nodes of type *t*:

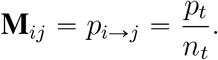

After constructing **M**, we finally compute the diffusion profile through power iteration as shown in Algorithm 1. The key equation is:

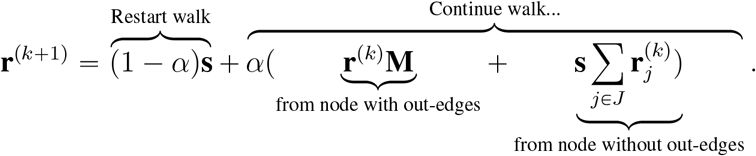

At each step, the random walker can restart its walk at the drug or disease node according to (1 – *α*)**s** or continue its walk. If the random walker continues its walk from a node with out-edges, then it jumps to an adjacent node according to *α*(**r**^(*k*)^**M**). If the random walker continues its walk from a node without out-edges (i.e. a sink node), then it restarts its walk according to 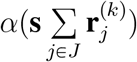 where *J* is the set of sink nodes in the graph. At every iteration, ∑_*i*_ **r**_*i*_ = 1.

Code for the power iteration implementation is available at github.com/snapstanford/multiscale-interactome. We use a tolerance of *∊* = 1 × 10^−6^.

**Algorithm 1.**
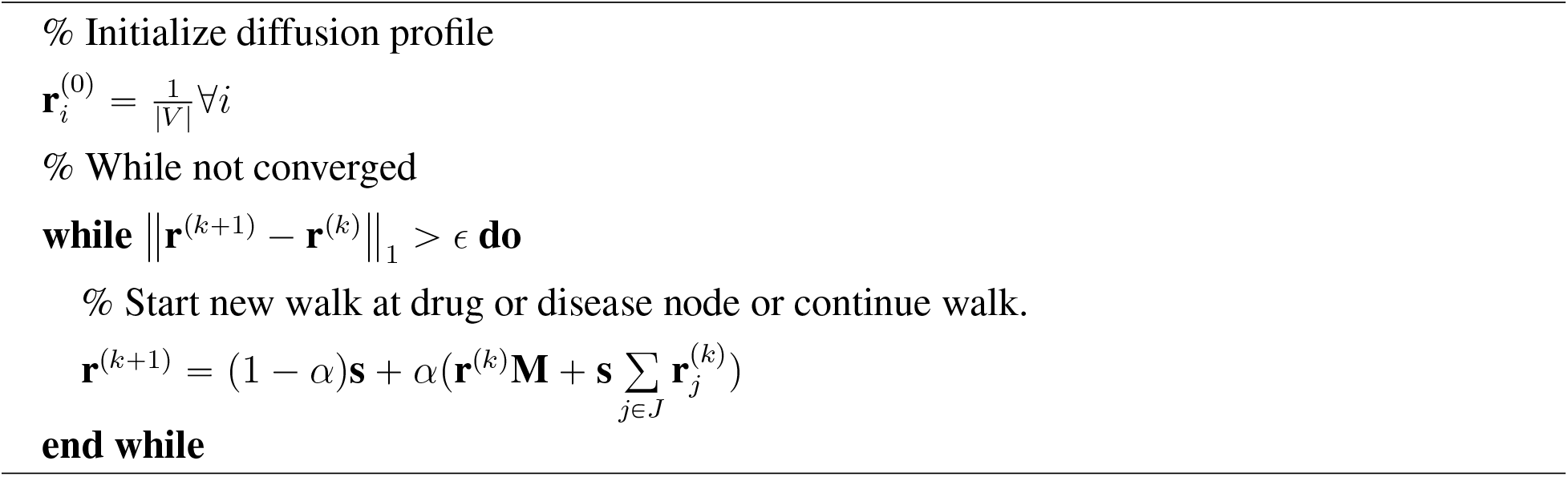
Diffusion profiles through power iteration

### Predicting what drugs will treat a given disease with diffusion profiles

For a drug to treat a disease, it must affect proteins and biological functions similar to those disrupted by the disease. The diffusion profiles of the drug **r**^(*c*)^ and the disease **r**^(*d*)^ encode the effect of the drug and the disease on proteins and biological functions. Therefore, comparing **r**^(*c*)^ and **r**^(*d*)^ allows us to predict what drugs treat a given disease.

For each drug and each disease, we compute the diffusion profile as described above. For each disease, we then rank-order the drugs most likely to treat the disease based on the similarity of the drug and disease diffusion profiles SIM(**r**^(*c*)^, **r**^(*d*)^) and a series of baseline methods.

We test five metrics of vector similarity:

1. L2 norm: 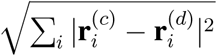,
2. L1 norm: 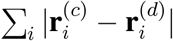,
3. Canberra distance: 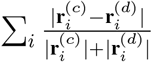,
4. Cosine similarity: 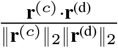,
5. Correlation distance: 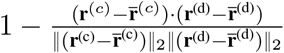.

We additionally test two proximity metrics. In particular, we consider the visitation frequency of the drug node *i* in the disease diffusion profile as: 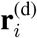. We also consider the visitation frequency of the drug node *i* in the disease diffusion profile multiplied by the visitation frequency of the disease node *j* in the drug diffusion profile: 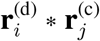.

### Baseline metrics to predict what drugs will treat a disease

To predict what drugs will treat a given disease, we consider baselines that measure (1) the overlap between drug targets and disease proteins, (2) the overlap between the functions of drug targets and disease proteins, and (3) the state-of-the-art proximity metric on a molecular-scale interactome. First, we compute the “protein overlap” baseline which we define as the Jaccard Similarity between the set of drug targets *T* and the set of disease proteins 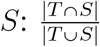. Second, we compute the “functional overlap” baseline which we define as SimIC which measures the semantic similarity between the GO terms *U* associated with the drug targets and the GO terms *V* associated with the disease proteins [96]. We tested 17 functional overlap baselines, of which this was the best performing (Methods: Baseline metrics of functional overlap between drug targets and disease proteins) (Supplementary Fig. 5). Third, we compute the state-of-the-art proximity metric on a molecular-scale interactome which is the closest distance metric in [10, 13]. Let *T* be the set of drug targets, *S* be the set of disease proteins, and *l*(*s, t*) be the shortest path length between nodes *s* and *t*. The state-of-the-art proximity metric first computes the “closest” distance 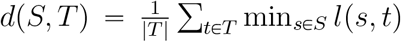 between *S* and *T*. Next, this distance is compared to a reference distance distribution which measures *d*(*S, T*) when *S* and *T* are randomly permuted to sets of proteins that match the size and degrees of the original disease proteins and drug targets in the network. Finally, the state-of-the-art proximity metric is computed by taking a z-score of *d*(*S, T*) with respect to the reference distribution: 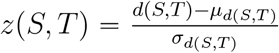.

### Baseline metrics of functional overlap between drug targets and disease proteins

We tested 17 baseline methods that predict what drugs treat a disease by considering the biological functions affected by drug targets and disease proteins (Supplementary Fig. 5).

First, we tested baseline methods that compare the functional overlap between drug targets and disease proteins. Let *U* and *V* be the sets of Gene Ontology (GO) terms associated with drug targets and disease proteins respectively. Let *U′* and *V′* be the multisets of GO terms associated with drug targets and disease proteins respectively. Let *U′′* and *V′′* be the sets of GO terms enriched among drug targets and disease proteins according to Gene Set Enrichment Analysis (GSEA) respectively [89, 97]. Note that in the multisets *U′* and *V′*, 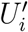 and 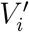 correspond to the number of occurrences of the *i*^th^ element in the multiset.

We measure the following baselines:

- The Jaccard Similarity or Intersection between the set of GO terms associated with the drug targets and the set of GO terms associated with the disease proteins: 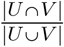 or |*U* ∩ *V*|
- The Jaccard Similarity or Intersection between the multiset of GO terms associated with the drug targets and the multiset of GO terms associated with the disease proteins: 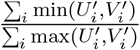 or ∑_*i*_ min 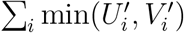
- The Jaccard Similarity or Intersection between the set of GO terms enriched among drug targets and the set of GO terms enriched among disease proteins according to Gene Set Enrichment Analysis [89, 97]: 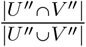 or |*U′′* ∩ *V′′*|
- The Z-scored Jaccard Similarity or Intersection between the set of GO terms associated with the drug targets and the set of GO terms associated with the disease proteins: 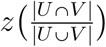 or *z*(|*U* ∩ *V*|)
- The Z-scored Jaccard Similarity or Intersection between the multisets of GO terms associated with the drug targets and the set of GO terms associated with the disease proteins: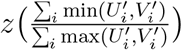 or 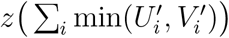

We compute reference distributions for z-scored metrics by following the approach in [10, 13]. Specifically, we randomly permute the set of disease proteins *S* and the set of drug targets *T* to sets of proteins that match the size and degrees of the original disease proteins and drug targets in the network. We then generate the GO sets and multisets that correspond to the permuted *S* and *T*, compute the relevant baseline metric, and repeat this for random permutations of *S* and *T* to generate a reference distribution. Finally, we compute a z-score by comparing the baseline metric for the true *S* and *T* to the reference distribution.

Second, we tested baseline methods that calculate the semantic similarity between the GO terms associated with the drug targets and those associated with the disease proteins [98]. Consider *U* and *V*, the sets of GO terms directly associated with drug targets and disease proteins respectively. Semantic similarity methods first define a similarity sim(*u, v*) between a GO term directly associated with drug targets *u* and a GO term directly associated with disease proteins *v*. The similarity of the sets *U* and *V* are subsequently calculated by aggregating across the similarities of pairwise GO terms *u* and *v*.

We used the following semantic similarity metrics as as they are among the most common and best-performing metrics in a variety of settings [98].

- The Resnik Similarity [99, 100] between *u* and *v* measures the information content of the most informative common ancestor between *u* and *v*. sim(*u, v*) = Resnik(*u, v*) = IC[MICA(*u, v*)]
  – Let *p*(*u*) be the fraction of proteins in the multiscale interactome that are associated with a GO term *u* or its descendants. The information content IC of term *u* is defined as IC(*u*) = − log[*p*(*u*)]. The Maximum Informative Common Ancestor (MICA) between two GO terms *u* and *v* is defined as 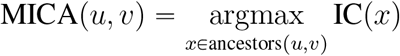.
- simIC [96] integrates both the information content of GO terms and the structural information of the GO hierarchy to determine the similarity between GO terms *u* and *v*: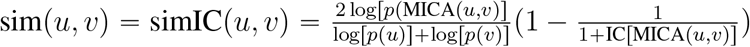
- simGIC [101] which considers the information content of all common ancestors of the GO terms directly associated with the drug targets *U* and the GO terms directly associated with the disease proteins 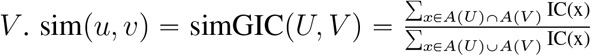.
  – Here, *A*(*X*) is the set of terms within *X* and all their ancestors in the GO hierarchy.

We aggregated the Resnik Similarity and simIC across *U* and *V* by using the average, maximum, and best match average approaches.

- Average: 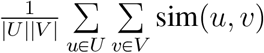
- Max: 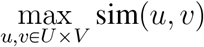
- Best Match Average [102]: 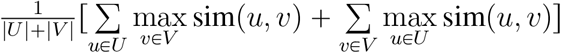

### Evaluating predictions of what drugs will treat a disease

We evaluate how effectively a model ranks the drugs that will treat a disease by using AUROC, Average Precision, and Recall@50. For each disease, a model produces a ranked list of drugs. We identify the drugs approved to treat the disease and, consistent with prior literature, assume that other drugs cannot treat the disease [11–14]. For each disease, we then compute the model’s AUROC, Average Precision, and Recall@50 values based on the ranked list of drugs. We report the model’s performance across diseases by reporting the median of the AUROC, the mean of the Average Precision, and the mean of the Recall@50 values across diseases.

To ensure robust results, we perform five-fold cross validation. We split the drugs into five folds and create training and held-out sets of the drugs and their corresponding indications. We compute the above evaluation metrics separately on the training and held-out sets. Ultimately, we report all performance metrics on the held-out set, averaged across folds (Figure 2b).

### Model selection and optimization of scalar weights

The diffusion profiles of each drug and disease depend on the scalar weights used to compute them *W* = {*w*_drug_, *w*_disease_, *w*_protein_, *w*_biological function_, *w*_higher-level biological function_, *w*_lower-level biological function_} and the probability *α* of continuing a walk. Similarly, how effectively diffusion profiles predict what drugs treat a given disease depends on the similarity metric used to compare drug and disease diffusion profiles. We optimize the prediction model across the scalar weights *W*, the probability of continuing a walk *α*, and the comparison metrics by performing a sweep and selecting the model with the highest median AUROC on the training set, averaged across folds.

After initial coarse explorations for each hyperparameter, we sweep across 486 combinations of hyperparameters sampled linearly within the following ranges *w*_drug_ ∈ [3, 9], *w*_disease_ ∈ [3, 9], *w*_protein_ ∈ [3, 9], *w*_higher-level biological function_ ∈ [1.5, 4.5], *w*_lower-level biological function_ ∈ [1.5, 4.5] *α* ∈ [0.85, 0.9] and set *w*_biological function_ = *w*_higher-level biological function_ + *w*_lower-level biological function_. We also sweep across the seven comparison metrics described above. We repeat this procedure for both the multiscale interactome and the molecular-scale interactome to identify the best diffusion-based model for both. The optimal weights for the molecular-scale interactome are *w*_drug_ = 4.88, *w*_disease_ = 6.83, *w*_protein_ = 3.21 with *α* = 0.854 and use the L1 norm to compare **r**^(*c*)^ and **r**^(*d*)^ (Figure 2c, Supplementary Note 1). The optimal weights for the multiscale interactome are *w*_drug_ =, *w*_disease_ = 3.54, *w*_protein_ = 4.40, *w*_higher-level biological function_ = 2.10, *w*_lower-level biological function_ = 4.49, *w*_biological function_ = 6.58 with *α* = 0.860 and use the correlation distance to compare **r**^(*c*)^ and **r**^(*d*)^ (Figure 2b, c). We utilize these optimal weights for the multiscale interactome for all subsequent sections. Optimized diffusion profiles are provided in Supplementary Data 10.

Additional information on selecting the edge weight ranges is provided as Supplementary Note 2.

### Evaluating predictions of what drugs will treat a disease by drug category

We analyze the multiscale interactome’s predictive performance across drug categories by using the Anatomical Therapeutic Chemical Classification (ATC) [103]. We map all drugs to their ATC class by using DrugBank’s XML database “full_database.xml” [30]. We use the second level of the ATC classification and only consider categories with at least 20 drugs. For the drugs in each ATC Level II category, we compute the rank of the drugs for the diseases they are approved to treat. We conduct this analysis twice, first to understand the overall performance of the best multiscale interactome model (Supplementary Fig. 6) and second to understand the differential performance of the best multiscale interactome model compared to the best molecular-scale interactome model using diffusion profiles (Figure 2c; Supplementary Fig. 7). The ATC classification for the drugs in our study is provided in Supplementary Data 7.

### Diffusion profiles identify proteins and biological functions related to treatment

For a given drug-disease pair, diffusion profiles identify the proteins and biological functions related to treatment. For each drug-disease pair, we select the top *k* proteins and biological functions in the drug diffusion profile and in the disease diffusion profile. To explain the relevance of these proteins and biological functions to treatment, we induce a subgraph on these nodes and remove any isolated components. We set *k* “ 10 for the case studies in Figures 2g, 2h, and 3f. We focus on these nodes since the nodes ranked most highly in the diffusion profiles have the highest propagated effect and are thus considered the most relevant to treatment. Additionally, these top nodes also capture a substantial fraction of the overall visitation frequency in the diffusion profile (i.e. about 50% for Figures 2g, 2h). We additionally include the rankings of the top 20 proteins and biological functions for each case study as Supplementary Fig. 16-18.

### Validation of diffusion profiles through gene expression signatures

To validate drug diffusion profiles, we compare drug diffusion profiles to the drug gene expression signatures present in the Broad Connectivity Map [47, 48] (Figure 2f).

We map drugs in the Broad Connectivity Map to DrugBank IDs using PubChem IDs, drug names, and the DrugBank “approved_drug_links.csv” and “drugbank_vocabulary.csv” files [30].

Drugs in the Broad Connectivity Map have multiple gene expression signatures based on the cell line, the drug dose, and the time of exposure. However, drugs only have a single diffusion profile. We thus only consider drugs where activity is consistent across cell lines and select a single representative gene expression signature for each drug. To accomplish this, we follow Broad Connectivity Map guidelines [47, 48] as described next. For drugs:

1. We only consider drugs with similar signatures across cell lines (an inter-cell connectivity score >= 0.4) and with activity across many cell lines (an aggregated transcriptional activity score >= 0.3).
2. We only consider drugs that are members of the “touchstone” dataset: the drugs that are the most well-annotated and systematically profiled across the Broad’s core cell lines at standardized conditions. The Broad Connectivity Map specifically recommends the “touchstone” dataset as a reference.

For gene expression signatures, we utilize the Level 5 Replicate Consensus Signatures provided by the Broad Connectivity Map. Each gene expression signature captures the z-scored change in expression of each gene across replicate experiments (“GSE92742_Broad_LINCS_Level5_COMPZ.MODZ_n473647x12328.gctx”). For these gene expression signatures:

1. We only consider genes whose expression is measured directly rather than inferred (i.e. “landmark” genes).
2. We only consider signatures that are highly reproducible and distinct (distil_cc_q75 >= 0.2 and (pct_self_rank_q25 <= 0.1).
3. We require that each signature be an “exemplar” signature for the drug as indicated by the Broad Connectivity Map (i.e. a highly reproducible, representative signature).
4. We require that each signature be sufficiently active (i.e. have a transcriptional activity score >= 0.35) and result from at least 3 replicates (distil_n_sample_thresh >= 3).
5. In cases where multiple signatures meet these criteria for a given drug, we select the signature with the highest transcriptional activity score.

The gene expression signatures we ultimately use for each drug are provided in Supplementary Data 8.

Finally, we compare the similarity of drugs based on their diffusion profiles and their gene expression signatures. We compare the similarity of drug diffusion profiles by the Canberra distance, multiplied by −1 so higher values indicate higher similarity. We compare the similarity of drug gene expression signatures based on the overlap in the 25 most upregulated genes *U* and 25 most downregulated genes *D*:

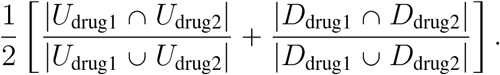

We use rank transformed gene expression signatures and diffusion profiles. We only allow the comparison of gene expression signatures that are in the same cell, with the same dose, and at the same exposure time. Ultimately, we measure the Spearman Correlation between the similarity of the drugs as described by the drug diffusion profiles and the similarity of the drugs as described the gene expression signatures.

### Compiling genetic variants that alter treatment

We compile genetic variants that alter treatment by using the Pharmacogenomics Knowledgebase (PharmGKB) [64]. PharmGKB is a goldstandard database mapping the effect of genetic variants on treatments. PharmGKB is manually curated from a range of sources, including the published literature, the Allele Frequency Database, the Anatomical Therapeutic Chemical Classification, ChEBI, ClinicalTrials.gov, dbSNP, Drug-Bank, the European Medicines Agency, Ensembl, FDA Drug Labels at DailyMed, GeneCard, HC-SC, HGNC, HMDB, HumanCyc Gene, LS-SNP, MedDRA, MeSH, NCBI Gene, NDF-RT, PMDA, PubChem Compound, RxNorm, SnoMed Clinical Terminology, and UniProt KB.

We use PharmGKB’s “Clinical Annotations” which detail how variants at the gene level alter treatments. PharmGKB’s “clinical ann metadata.tsv” file provides triplets of drugs, diseases, and genetic variants known to alter treatment. Treatment alteration occurs when a genetic variant alters the efficacy, dosage, metabolism, or pharmacokinetics of treatment or otherwise causes toxicity or an adverse drug reaction. We map genes to their Entrez ID using HUGO, drugs to their DrugBank ID using PharmGKB’s “drugs.tsv” and “chemicals.tsv” files, and diseases to their UMLS CUIDs by using PharmGKB’s “phenotypes.tsv” file. To ensure consistency with the approved drug-disease pairs we previously compiled, we only consider (drug, disease, gene) triplets in which the drug and disease are part of an FDA-approved treatment. Ultimately, we obtain 1,223 drug-disease-gene triplets with 201 drugs, 94 diseases, and 455 genes. All drug-disease-gene triplets are provided in Supplementary Data 9.

### Computing treatment importance of a gene based on diffusion profiles

We define the treatment importance (TI) of gene *i* as the product of the visitation frequency of the corresponding protein in the drug and disease diffusion profiles. For a treatment composed of drug compound *c* and disease *d*, the treatment importance of gene *i* is:

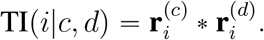

We define the treatment importance percentile as the percentile rank of TI(*i*|*c, d*) compared to all other genes for the same drug and disease. Intuitively, gene *i* is important to a treatment if the corresponding protein is frequently visited in both the drug and disease diffusion profiles.

### Comparing treatment importance of treatment altering genetic mutations vs other genetic mutations

We compare the treatment importance of genes known to alter a treatment with the treatment importance of other genes (Figure 3b). In particular, we compare the set of (drug, disease, gene) triplets where the gene is known to alter the drug-disease treatment with an equivalently sized set of (drug, disease, gene) triplets where the gene is not known to alter treatment. We construct the latter set by sampling drugs, diseases, and genes uniformly at random that are not known to alter treatment from PharmGKB [64]. The drugs and diseases in all triplets correspond to approved drug-disease pairs. Thereby, we construct a distribution of the treatment importance for “treatment altering genes” and a distribution of the treatment importance for “other genes” (Figure 3b).

### Predicting genes that alter a treatment based on treatment importance

We evaluate the ability of treatment importance to predict the genes that will alter a given treatment (Figure 3c). For each (drug, disease, gene) triplet, we use the treatment importance of the gene TI(*i*|*c, d*) to predict whether the gene alters treatment or not for that drug-disease pair (i.e. binary classification). We use the set of positive and negative (drug, disease, gene) triplets constructed previously (see Methods: Comparing treatment importance of treatment altering genetic mutations vs other genetic mutations). We assess performance using AUROC and Average Precision (Figure 3c).

### Comparing treatment importance of genes that alter one drug indicated to treat a disease but not another

We analyze how often a gene has a higher treatment importance in the treatments it alters than in those it does not alter (Figure 3e).

Formally, let *i* be a gene. Consider a triplet (*d, c*_altered_, *c*_unaltered_) of a disease *d*, a drug *c*_altered_ approved to treat the disease whose treatment is altered due to a mutation in *i*, and a drug *c*_unaltered_ approved to treat the disease whose treatment is not altered due to a mutation in *i*. Let *n*_triplets_ be the total number of such triplets for gene *i*. For each gene *i*, we measure the fraction *f* of triplets (*d, c*_altered_, *c*_unaltered_) for which the treatment importance of *i* is higher in the (*c*_altered_*, d*) treatment than in the (*c*_unaltered_, *d*) treatment, as shown below. We only consider genes for which *n*_triplets_ *≥* 100.

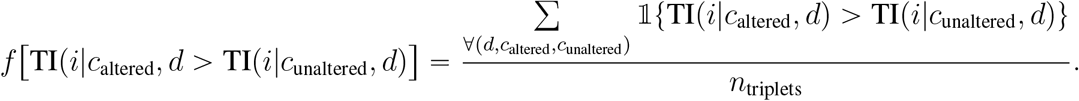

### Analyzing whether distant proteins can have common biological functions

We analyzed whether two proteins can be more distant than expected by random chance in a physical protein-protein interaction (PPI) network yet affect the same function (Supplementary Fig. 2). To run this analysis, we first compute the set of all protein pairs that are both present in the protein-protein interaction network described previously (Methods: Protein–protein interactions) and are also associated with a common biological function. We only consider direct associations of proteins to biological functions (i.e. we do not propagate associations up the GO hierarchy) in order to ensure that shared biological functions are specific and not generic (i.e. shared associations with the GO term ‘Biological Process’).

For each protein pair with a common biological function, we then:

1. Compute the shortest path distance in the PPI network between these two proteins.
2. Construct a reference distribution of shortest paths for these two protein pairs by following the approach in [10, 13]. Specifically, we randomly sample other proteins in the network with similar degree to the original proteins and measure the shortest path distance. These randomly sampled proteins do *not* necessarily share a common biological function.
3. Using the true shortest path distance between the proteins and the random reference distribution, we compute a z-score. The z-score captures whether the proteins with a shared function are closer or further away than expected by random chance in the PPI network.

### Construction of alternative multiscale interactomes that explicitly represent cells, tissues, and organs

We constructed three alternative multiscale interactomes which explicitly represent cells, tissues, and organs. In these alternative multiscale interactomes, the nodes and edges in the original multiscale interactome are all present. Additionally, (1) human cells, tissues, and organs are added as additional nodes; (2) edges between these cell, tissue, and organ nodes are added according to relationships defined in established anatomical ontologies; and (3) edges between GO biological function nodes and cell, tissue, and organ nodes are added according to relationships provided in Gene Ontology Plus (GO Plus) [104]. GO Plus maintains a curated set of relationships between the biological functions in GO and the cell, tissue, and organ nodes present in two key anatomical ontologies: Uberon and the Cell Ontology. We thus constructed three alternative multiscale interactomes incorporating human subsets of Uberon, the Cell Ontology, and both Uberon and the Cell Ontology.

1. *Multiscale Interactome + Uberon*: Uberon is an ontology covering anatomical structures in animals [105, 106]. Uberon nodes include tissues (i.e. cardiac muscle tissue UBERON:0001133), organs (i.e. heart UBERON:0000948), and organ systems (i.e. cardiovascular system UBERON:0004535). We utilized GO Plus (i.e. “go-plus.owl”) to link GO biological function nodes present in our original network to Uberon nodes present in a human-specific subset of Uberon (i.e. “subsets/human-view.obo”). Edges between Uberon nodes, which encode anatomical relationships, were also added according to “subsets/human-view.obo”.
2. *Multiscale Interactome + Cell Ontology*: The Cell Ontology is an ontology for the representation of in vivo cell types [107, 108]. Nodes consist primarily of cell types and their hierarchical relationships (i.e. epithelial cell CL:0000066, epithelial cell of pancreas CL:0000083, pancreatic A cell CL:0000171). We utilized a human-specific subset of the Cell Ontology previously prepared by the Human Cell Atlas Ontology [109]. We utilized GO Plus to link GO biological function nodes in our original network to Cell Ontology terms and the Cell Ontology (i.e. “cl-basic.obo”) to link Cell Ontology terms with one another.
3. *Multiscale Interactome + Uberon + Cell Ontology*: The “Multiscale Interactome + Uberon + Cell Ontology” network contains all nodes and edges present in our original network as well as nodes and edges added via GO Plus, Uberon, and Cell Ontology as described above.

### Prediction of what drugs treat a given disease in alternative multiscale interactomes

We evaluate the ability of diffusion profiles to predict what drugs treat a given disease in the alternative multiscale interactomes (see Methods: Construction of alternative multiscale interactomes that explicitly represent cells, tissues, and organs). Given the presence of new node types, we modify the edge weight hyperparameters used in the calculation of diffusion profiles. We then sweep over the full set of edge weight hyperparameters according to the broad hyperparameter sweep described in Supplementary Note 2, in which we sample 586 combinations of hyperparameters sampled linearly in the range [1, 100]. The new sets of edge weight hyperparameters and their optimal values are present below:

1. *Multiscale Interactome + Uberon*: The optimal weights for “Multiscale Interactome + Uberon” are *w*_drug_ = 55.2, *w*_disease_ = 27.3, *w*_protein_ = 76.8, *w*_biological function_ = 66.1, *w*_uberon_ = 82.2, *w*_higher-level biological function or uberon_ = 67.1, *w*_lower-level biological function or uberon_ = 45.7 with *α* = 0.76 and use the correlation distance to compare **r**^(*c*)^ and **r**^(*d*)^.
2. *Multiscale Interactome + Cell Ontology*: The optimal weights for “Multiscale Interactome + Cell Ontology” are *w*_drug_ = 39.0, *w*_disease_ = 17.1, *w*_protein_ = 72.4, *w*_biological function_ = 60.0, *w*_cell ontology_ = 23.1, *w*_higher-level biological function or cell ontology_ = 25.7, *w*_lower-level biological function or cell ontology_ = 22.8 with *α* = 0.83 and use the correlation distance to compare **r**^(*c*)^ and **r**^(*d*)^.
3. *Multiscale Interactome + Uberon + Cell Ontology*: The optimal weights for “Multiscale Interactome + Uberon + Cell Ontology” are *w*_drug_ = 60.2, *w*_disease_ = 12.8, *w*_protein_ = 42.3, *w*_biological function_ = 78.4, *w*_uberon_ = 70.0, *w*_cell ontology_ = 91.7, *w*_higher-level biological function or uberon or cell ontology_ = 26.7, *w*_lower-level biological function or uberon or cell ontology_ = 76.1 with *α* = 0.82 and use the correlation distance to compare **r**^(*c*)^ and **r**^(*d*)^.

