## Supplementary Information for "Identification of disease treatment mechanisms through the multiscale interactome"

This PDF file includes:

Supplementary Notes 1 to 5

Supplementary Figures 1 to 18

Other supporting material for this manuscript includes the following. Data and code are available at github.com/snap-stanford/multiscale-interactome.

Supplementary dataset of interactions between drugs and proteins

Supplementary dataset of interactions between diseases and proteins

Supplementary dataset of interactions between proteins and proteins

Supplementary dataset of interactions between proteins and biological functions

Supplementary dataset of hierarchy of interactions between biological functions

Supplementary dataset of approved drug-disease pairs

Supplementary dataset of drug classes according to the Anatomical Therapeutic Chemical Classification

Supplementary dataset of selected gene expression signatures from the Broad Connectivity Map

Supplementary dataset of genetic mutations that alter treatment from PharmGKB

Supplementary dataset of optimized diffusion profiles for drugs and diseases

#### Supplementary Note 1 Predictive power of diffusion profiles on a molecular-scale interactome.

We find that diffusion profiles improve prediction of what drugs treat a disease, even when computed on a molecular-scale interactome. We compare optimized diffusion profiles on the molecular-scale interactome to a state of the art proximity metric[1,2] (Methods). We find that diffusion profiles improve prediction relative to proximity (AUROC = 0.674 vs. 0.620, +12.3%; Average Precision = 0.078 vs. 0.065, +20.0%; Recall@50 = 0.317 vs. 0.264, +20.0%). All evaluation metrics are reported on a held-out set, averaged across five-fold cross validation (Methods).

#### Supplementary Note 2 Selecting ranges for optimization of scalar weights

The multiscale interactome utilizes optimized edge weights to propagate the effects of drugs and diseases to proteins and biological functions. To optimize these edge weights, we conduct a hyperparameter sweep and allow each hyperparameter to vary over a prespecified range. In our initial sweep, we conduct coarse explorations for each hyperparameter in the range [1, 10] and then conduct a finer sweep in the range [3, 9]. Notice that in the calculation of a diffusion profile, the relative values of two hyperparameters dictate the likelihood that the random walker visits one node type rather than another. For example, if $w_{\text{biological function}} = 9$ and $w_{\text{protein}} = 3$, then a random walker adjacent to both a biological function node and a protein node is three times more likely to visit the biological function node than the protein node. If instead $w_{\text{biological function}} = 3$ and $w_{\text{protein}} = 1$, then the random walker is still three times more likely to visit the biological function node than the protein node. In other words, the relative values between the weights affect the propagation but the absolute value of each particular weight does not. By setting the range of each hyperparameter to [3, 9], we thus allow each node type to be jumped to as much as 3X more or less often than other node types.

To ensure that our initial hyperparameter sweep was not too narrow, we conduct a broader hyperparameter sweep and test 560 combinations of edge weight hyperparameters sampled linearly in the range [1, 100] and $\alpha \in [0.667, 1.0]$. Here, each node type can be jumped to as much as 100X more or less often than other node types depending on the sampled weights. The optimal model identified by this expanded hyperparameter sweep did not outperform the model previously described (Methods: Model selection and optimization of scalar weights) in predicting what drug treats a given disease (AUROC: 0.706 vs 0.705; Average Precision: 0.092 vs 0.091; Recall@50: 0.335 vs 0.347). Therefore, we empirically find that our initial hyperparameter range was appropriate and not too narrow.

#### Supplementary Note 3 Identifying proteins and biological functions relevant to treatment

**The multiscale interactome identifies proteins and biological functions relevant to treatment from a large number of possibilities.** To explain treatment, the multiscale interactome identifies a subgraph of relevant proteins and biological functions. This subgraph explains how drug-protein, disease-protein, protein-protein, protein-function, and function-function edges compose in a manner that explains treatment. Critically, our case studies describe treatments that cannot be explained by a single protein node, biological function node, protein-protein edge, or protein-function edge. Instead, treatment results from the complex interactions of multiple proteins and biological functions, and a subgraph is necessary to illustrate how these interactions compose to explain treatment. Additionally, note that the multiscale interactome is not meant to identify new protein-function edges or new edges generally. The protein-function edges and other edges in our network are already known. Instead, our model is meant to identify how existing interactions between drugs, diseases, proteins, and biological functions compose to explain treatment.

Moreover, note that the multiscale interactome identifies relevant proteins and biological functions from a large array of possibilities. In each case study, we identify 18 proteins and biological functions relevant to treatment from a network of 17,660 proteins and 9,798 biological functions. Notice that there are $1.22 \times 10^{64}$ potential choices of proteins and biological functions (27,458 choose 18). (For comparison, note there are only about $7 \times 10^{22}$ stars in the entire universe.) From this large array of possibilities, diffusion profiles identify a subgraph of proteins and functions that are relevant to treatment. We validate the subgraphs by comparison with externally validated treatment mechanisms. Notably, the knowledge from these externally validated treatment mechanisms is never encoded in our model. Instead, diffusion profiles identify relevant proteins and biological functions from a large array of possibilities. This is the key explanatory benefit of our approach.

**Alternative approaches to identify biological functions relevant to treatment.** We compared the ability of the multiscale interactome to identify biological functions relevant to treatment with three alternative approaches: selecting GO terms associated with both drug targets and disease proteins, selecting GO terms enriched among both drug targets and disease proteins according to Gene Set Enrichment Analysis[3], and selecting GO terms on shortest paths between drug targets and disease proteins. First, we discuss conceptual limitations of these alternative approaches. Second, we compare these approaches with the multiscale interactome directly for the case studies presented as Figure 2g and Figure 2h.

**Conceptual Issues with Alternative Approaches:** Conceptually, these alternative approaches are limited in ways that apply generally regardless of the case study utilized. Selecting common GO terms between drug targets and disease proteins will necessarily prioritize the most popular GO terms that the most proteins connect to (i.e. Biological Process, Cellular Process, Metabolic Process) which are too generic to yield meaningful insight into treatment. Selecting GO terms enriched among drug targets and disease proteins struggles in cases where there are few drug targets or disease proteins. In these cases, few GO terms are enriched and there are few or no common enriched terms, thus yielding little insight. Identifying GO terms that occur on the shortest paths between drug targets and disease proteins fails in cases where the shortest paths between drug targets and disease proteins do not include GO terms. In cases where shortest paths do include the relevant GO terms, diffusion profiles are likely to identify these GO terms anyways since nodes on the shortest paths between drug targets and disease proteins are frequently visited in diffusion profiles. Finally, all three alternative approaches only generate *lists* of GO terms. By contrast, diffusion profiles identify *subgraphs* which demonstrate how complex interactions between drugs, diseases, proteins, and biological functions can help explain treatment. Lists of GO terms can at most identify relevant functions but cannot shed light into the drug-protein, disease-protein, protein-protein, protein-function, or function-function interactions that are relevant to treatment. Such complex interactions are important for describing treatment[1,2,4–7].

**Example: Treatment of Cryopyrin-Associated Periodic Syndromes by Anakinra.** First, consider the treatment of Cryopyrin-Associated Periodic Syndromes by Anakinra (Figure 2h). Cryopyrin-Associated Periodic Syndromes are characterized by immune-mediated inflammation via the Interleukin-1 beta signaling pathway[8]. Anakinra treats Cryopyrin-Associated Periodic Syndromes by binding to a regulator of the Interleukin-1 beta signaling pathway and thus preventing excess inflammation[9,10]. Diffusion profiles capture Anakinra's treatment of Cryopyrin-Associated Periodic Syndromes by affecting immune-mediated inflammation via the Interleukin-1 beta signaling pathway (Supplementary Figure 14a). Diffusion profiles identify 14 GO terms, 10 of which describe treatment: "inflammatory response," "response to interleukin-1," "IL-1 mediated signaling pathway," "cytokine-mediated signaling pathway," "negative regulation of IL-1 mediated signaling pathway," "regulation of IL-1 mediated signaling pathway," "negative regulation of inflammatory response," "cellular response to IL-1," "cellular response to cytokine stimulus," and "response to cytokine." Diffusion profiles thus capture the key biological functions underlying treatment: regulation of immune-mediated inflammation via the Interleukin-1 beta signaling pathway.

By contrast, none of the alternative approaches capture the key biological function underlying treatment: regulation of immune-mediated inflammation via the Interleukin-1 beta signaling pathway. First if we consider GO terms that are associated with both Anakinra's targets and Cryopyrin-Associated Periodic Syndrome's genes, we find nine such GO terms (Supplementary Figure 14b). However, these GO terms broadly miss the effect of Anakinra on immune-mediated inflammation and the Interleukin-1 beta signaling pathway. Indeed, eight of these nine GO terms are generic and not useful to describing treatment (i.e. "biological process," "regulation of cellular process," "response to chemical," "biological regulation," "cellular process," "regulation of biological process," "response to organic substance," "response to stimulus"). Second if we consider GO terms enriched among both Anakinra's targets and Cryopyrin Associated Periodic Syndrome's genes, we find that there are *no* common GSEA GO terms (Supplementary Figure 14c). Finally if we consider GO terms on shortest paths between Anakinra's targets Cryopyrin Associated Periodic Syndrome's genes, we find that there are no GO terms on the shortest paths between Anakinra and Cryopyrin Associated Periodic Syndromes (Supplementary Figure 14d). Therefore, the three alternative approaches miss the effect of Anakinra on immune-mediated inflammation and the Interleukin-1 beta signaling pathway. Diffusion profiles thus identify the biological functions related to the treatment of Cryopyrin-Associated Periodic Syndromes by Anakinra more effectively than alternative approaches.

**Example: Treatment of Hyperlipoproteinemia Type III by Rosuvastatin.** Next, consider the treatment of Hyperlipoproteinemia Type III by Rosuvastatin. Hyperlipoproteinemia is a disease characterized by abnormal levels of cholesterol in the blood[11–15]. Rosuvastatin treats Hyperlipoproteinemia Type III by affecting various biological functions related to cholesterol biosynthesis and metabolism[10,16,17]. Diffusion profiles successfully identify the cholesterol biosynthesis functions relevant to treatment of Hyperlipoproteinemia Type III's by Rosuvastatin (Supplementary Figure 13a). Diffusion profiles identify 12 GO terms, 5 of which describe cholesterol biosynthesis (i.e. "regulation of cholesterol biosynthetic process," "positive regulation of cholesterol biosynthetic process," "negative regulation of cholesterol biosynthetic process," "cholesterol biosynthetic process," "cholesterol homeostasis"). An additional 3 GO describe triglyceride metabolism, a function dysregulated in Hyperlipoproteinemia Type III[11–15] (i.e. "positive regulation of triglyceride catabolic process," "triglyceride metabolic process," "triglyceride homeostasis"). Diffusion profiles thus capture key biological functions relevant to treatment.

By contrast, two of the three alternative approaches are less effective than diffusion profiles in identifying biological functions that are both specific and relevant to treatment. First, if we consider

GO terms enriched among both Rosuvastatin's target and Hyperlipoproteinemia Type III's genes, we find no common enriched GO terms, thus yielding no insight into treatment (Supplementary Figure 13c). Second, if we consider GO terms that are associated with both Rosuvastatin's target and Hyperlipoproteinemia Type III's genes, only 1 of 19 identified GO terms is specific and relevant (Supplementary Figure 13b). The remaining GO terms are generic (i.e. "cellular component organization", "macromolecular complex assembly") and thus not particularly insightful. By contrast, 8 of the 12 GO terms identified by diffusion profiles are specific and relevant. Moreover, the single relevant GO term identified via this alternative approach (i.e. "cholesterol metabolic process") is more generic than the five cholesterol-related GO terms identified via diffusion profiles (i.e. "regulation of cholesterol biosynthetic process," "positive regulation of cholesterol biosynthetic process," "negative regulation of cholesterol biosynthetic process," "cholesterol biosynthetic process," "cholesterol homeostasis"). Indeed in the Gene Ontology, metabolic processes define *broad* functions that can be altered in many ways. Two *specific* ways to alter metabolic processes are effects on biosynthesis and biosynthesis regulation[18,19], the functions identified by diffusion profiles. Ultimately, common enriched GO terms and common associated GO terms are less effective than diffusion profiles in identifying biological functions both specific and relevant to treatment.

The third alternative approach, considering GO terms on shortest paths between Rosuvastatin and Hyperlipoproteinemia Type III, performs comparably to diffusion profiles, identifying 4 GO terms related to cholesterol biosynthesis (i.e. "cholesterol metabolic process," "negative regulation of cholesterol biosynthetic process," "cholesterol biosynthetic process," and "cholesterol homeostasis") (Supplementary Figure 13d). Three of these GO terms overlap with the biological functions identified by diffusion profiles. Here, the comparable performance between diffusion profiles and GO terms on shortest paths makes sense. In cases where relevant biological functions are on shortest paths between drug targets and disease proteins, diffusion profiles will identify these GO terms as they will likely be frequently visited in the drug and disease diffusion profiles. In other treatments, however, the shortest paths between drug targets and disease proteins may not include GO terms. For such treatments, diffusion profiles will provide functional insight not provided by simply identifying GO terms on shortest paths.

Diffusion profiles capture the cholesterol biosynthesis functions relevant to treatment and thus offer an effective approach to identifying the biological functions relevant to treatment.

### Supplementary Note 4 The multiscale nature of our network and alternative multiscale interactomes

The multiscale interactome contains both physical interactions between proteins and a hierarchy of biological functions. We consider the network "multiscale" for two reasons. First, biological functions are often defined at the level of molecules (i.e. DNA Demethylation GO:0080111), cells (i.e. Mitotic Cell Cycle GO:0000278), tissues (i.e. Muscle Atrophy GO:0014889), organ systems (i.e. Activation of Innate Immune Response GO:0002218), and the whole organism (i.e. Anatomical Structure Development GO:0048856)[20,21]. Second, the hierarchical relationships between biological functions contains a "detailed implicit ontology of anatomical structures"[22]. This point is made clearer in "GO-Plus"[19] which links biological functions to the cells, tissues, and organs in Uberon[22,23] and the Cell Ontology[24,25] (two anatomical ontologies) that these biological functions affect. For these reasons, we consider the network we constructed "multiscale."

We acknowledge that alternative interpretations of "multiscale" may wish that cells, tissues, and organs be explicitly included as nodes in our network along with the anatomical relationships between them. To address this potential interpretation, we construct alternative multiscale interactomes that explicitly represent cells, tissues, organs and the relationships between them (Supplementary Figure 8, Methods). Specifically, we created three alternative multiscale interactomes that incorporate human-specific subsets of Uberon[22,23], the Cell Ontology[24,25], or both. In these alternative multiscale interactomes, our initial network is expanded such that (1) human cells, tissues, and organs are added as additional nodes, (2) these cells, tissues, and organs have edges between them according to the anatomical hierarchies defined in human subsets of Uberon and the Cell Ontology, and (3) the biological functions in GO are linked to the new cell, tissue, and organ nodes that they affect according to Gene Ontology Plus (GO Plus)[19]. All three of the alternative multiscale interactomes outperform molecular-scale interactomes when predicting what drug treats a disease. The best alternative multiscale interactome (multiscale interactome + Uberon) outperforms a molecular-scale interactome by 15.3% in AUROC (0.715 vs. 0.620), 40.0% in Average Precision (0.091 vs. 0.065), and 29.9% in Recall@50 (0.343 vs. 0.264) (Supplementary Figure 8). In these alternative multiscale interactomes, our model is still able to effectively predict what drugs treat a disease.

### Supplementary Note 5 Variants in network construction and edge weighting

We test whether numerous variants to the edge weighting and network construction approach described in Methods would improve the ability of our model to predict what drugs treats a given disease.

1. The multiscale interactome with all GO biological function terms vs. the multiscale interactome with only biological function terms associated with at least one drug target or disease protein (Supplementary Figure 9; Methods: Biological function – biological function interactions).

2. The multiscale interactome when all disease-protein edges are treated equivalently (i.e. $w_{\text{protein}}$, $w_{\text{disease}}$) vs. the multiscale interactome when different disease-protein edge types receive different hyperparameter edge weights (i.e. $w_{\text{disease}\rightarrow\text{protein via genomic alteration}}$, $w_{\text{disease}\rightarrow\text{protein via altered expression}}$, $w_{\text{disease}\rightarrow\text{protein via post-translational modification}}$, $w_{\text{protein}\rightarrow\text{disease via genomic alteration}}$, $w_{\text{protein}\rightarrow\text{disease via gene expression}}$, $w_{\text{protein}\rightarrow\text{disease via post-translational modification}}$) (Supplementary Figure 11).

3. The multiscale interactome when all biological function – biological function edges are treated equivalently (i.e. $w_{\text{biological function}}$, $w_{\text{higher-level biological function}}$, $w_{\text{lower-level biological function}}$) vs. when they are differentiated based on the type of GO edge (i.e. $w_{\text{biological function}}$, $w_{\text{higher-level bf via 'is a'}}$, $w_{\text{lower-level bf via 'is a'}}$, $w_{\text{higher-level bf via 'part of'}}$, $w_{\text{lower-level bf via 'part of'}}$, $w_{\text{higher-level bf via 'regulates'}}$, $w_{\text{lower-level bf via 'regulates'}}$, $w_{\text{higher-level bf via 'positively regulates'}}$, $w_{\text{lower-level bf via 'positively regulates'}}$, $w_{\text{higher-level bf via 'negatively regulates'}}$, $w_{\text{lower-level bf via 'negatively regulates'}}$; "bf" denotes biological function) (Supplementary Figure 10).

For each variant, we conduct the broad hyperparameter sweep described in Supplementary Note 2. Hyperparameters not mentioned above are defined as in the original model (Methods). Ultimately, we use these variants to predict what drug treats a disease according to the approach previously described (Methods). All optimal models used the correlation distance to compare the drug and disease diffusion profiles. The described variants did not substantially outperform the original model in predicting what drugs treat a given disease (Supplementary Figure 9, Supplementary Figure 11, Supplementary Figure 10).

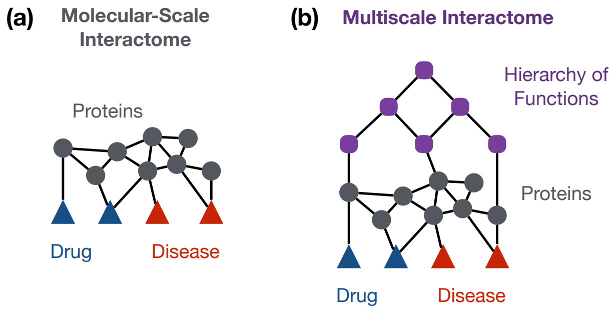


**Supplementary Figure 1: The multiscale interactome incorporates both physical interactions between proteins and a hierarchy of biological functions.** **(a)** Molecular scale interactomes model drug-disease treatment by primarily considering physical interactions (i.e. drug–protein, disease–protein, and protein–protein edges). **(b)** The multiscale interactome models drug-disease treatment by considering both physical interactions (i.e. drug–protein, disease–protein, protein–protein edges) and a hierarchy of biological functions (i.e. protein–biological function, biological function – higher-level biological function, and biological function – lower-level biological function edges). By modeling both physical interactions between proteins and a hierarchy of biological functions, the multiscale interactome integrates physical and functional relationships in the modeling of drug-disease treatment. The detailed construction of both networks is provided in Methods.

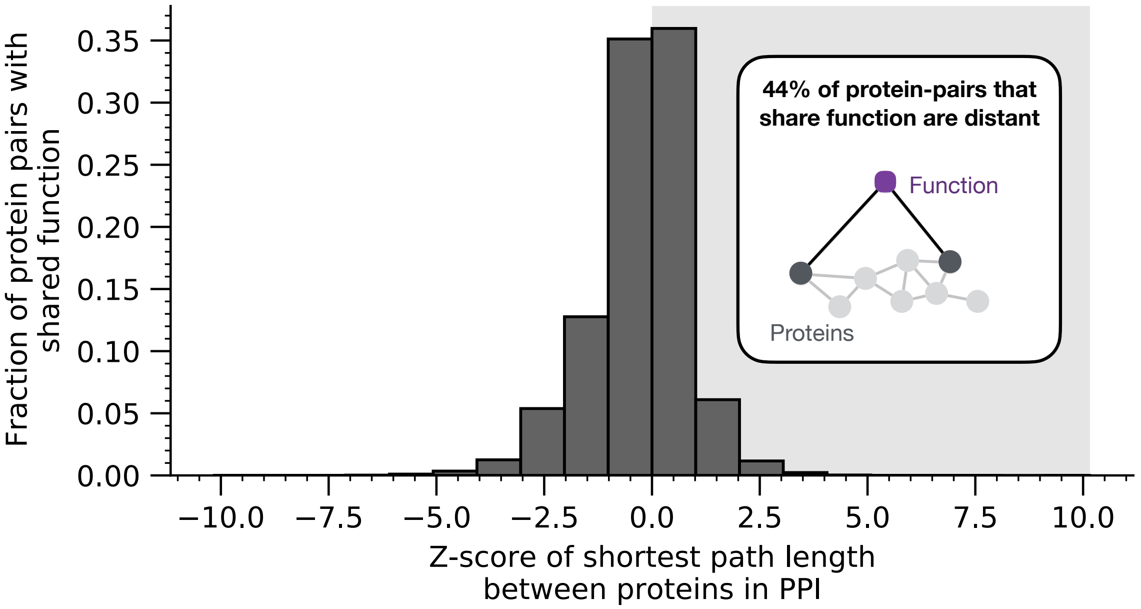


**Supplementary Figure 2: Biological functions capture critical relationships between proteins that physical PPI networks alone do not capture.** 44% of protein-protein pairs that affect the same biological function are more distant in protein-protein interaction (PPI) network than is expected by random chance. Therefore, proteins may be physically distant yet affect the same function, and biological functions capture critical relationships between proteins that physical protein-protein interaction networks alone do not capture. The shortest path length between each protein-pair in the PPI network that shares a function was compared to a reference distribution of shortest path lengths between protein pairs of similar degree (Methods).

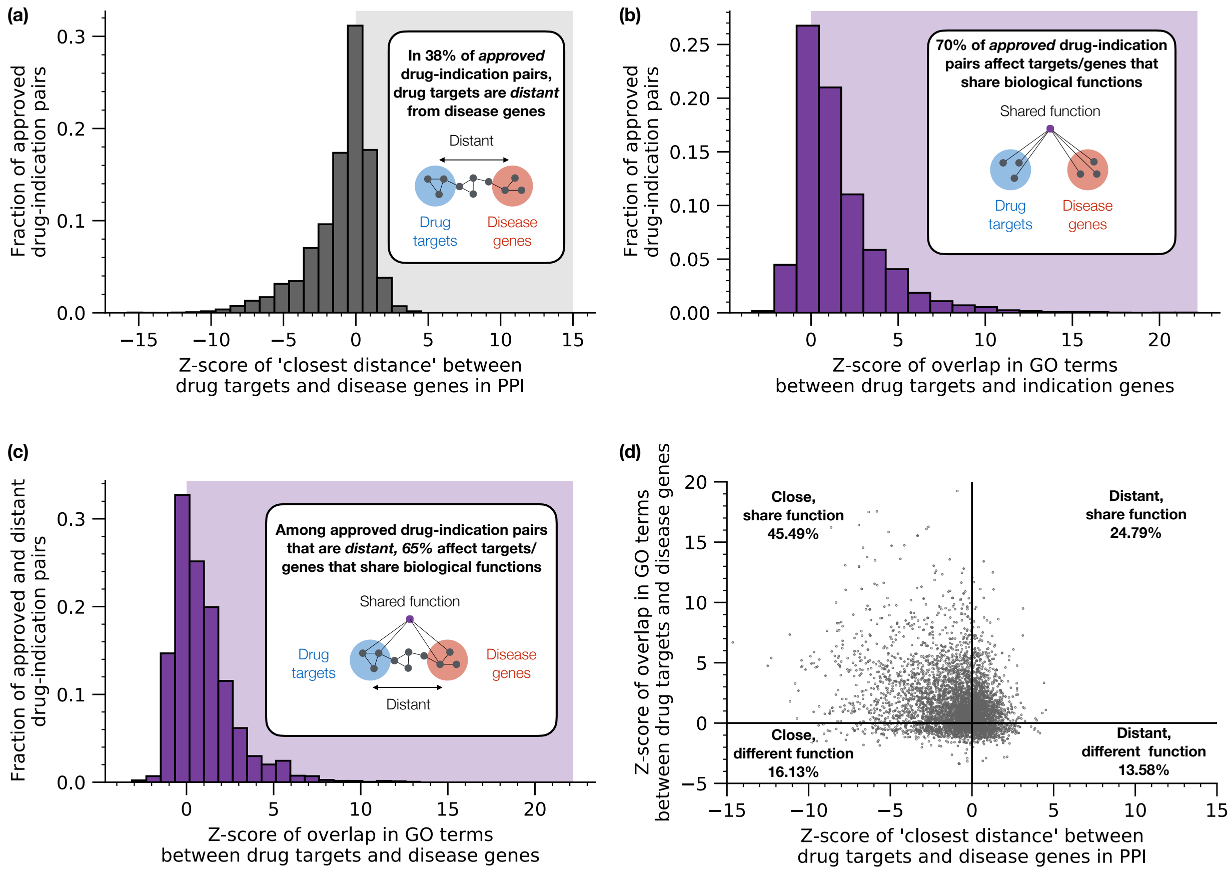


**Supplementary Figure 3: Drugs may treat diseases by affecting the same functions, even when drug targets are far from disease genes.** **(a)** In 38% of approved drug-indication pairs, drug targets are farther from disease genes than is expected by random chance. Distance between drug targets and disease genes is calculated according to a state-of-the-art proximity metric[1,2] (Methods). **(b)** In 70% of approved drug-indication pairs, drug targets affect the same biological functions as disease genes more often than is expected by random chance, suggesting that biological functions are broadly important in modeling treatment. The functional overlap metric used is the Z scored intersection of GO term multisets (Methods). **(c)** Among *distant* drug-indication pairs, 65% affect the same biological functions more often than is expected by random chance, suggesting that biological functions offer a powerful and complementary approach to explaining drug treatment. **(d)** Drugs treat diseases by either targeting proteins close to the disease genes (16.13% of approved drug-indication pairs), targeting proteins that affect the same biological functions as the disease genes (24.79%), or both (45.49%). Molecular-scale interactome approaches can only model proximity-based treatments. By integrating both physical interactions and biological functions, the multiscale interactome can model treatments that harness physical proximity, functional overlap, or both.

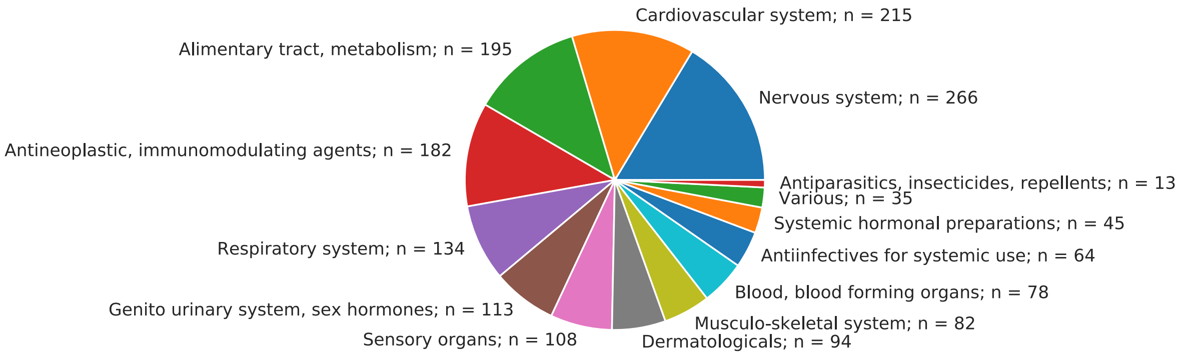


**Supplementary Figure 4: The multiscale interactome models treatments across human anatomy.** The multiscale interactome includes approved drug-disease treatments across human anatomy as represented by the Anatomical Therapeutic Chemical Classification (ATC). Each drug is mapped to its ATC Level I class. Numbers represent the number of unique drugs per class.

| Method | AUROC | Avg Prec | Rec@50 |
|---|---|---|---|
| Jaccard Similarity of GSEA GO Term Sets | 0.500 | 0.024 | 0.219 |
| Intersection of GSEA GO Term Sets | 0.500 | 0.022 | 0.217 |
| Jaccard Similarity of GO Term Sets | 0.564 | 0.048 | 0.235 |
| Intersection of GO Term Sets | 0.558 | 0.045 | 0.213 |
| Jaccard Similarity of GO Term Multisets | 0.560 | 0.041 | 0.216 |
| Intersection of GO Term Multisets | 0.554 | 0.040 | 0.208 |
| Z Scored Jaccard Similarity of GO Term Sets | 0.533 | 0.039 | 0.211 |
| Z Scored Intersection of GO Term Sets | 0.551 | 0.040 | 0.207 |
| Z Scored Jaccard Similarity of GO Term Multisets | 0.533 | 0.035 | 0.205 |
| Z Scored Intersection of GO Term Multisets | 0.553 | 0.039 | 0.196 |
| Max of Resnik Similarity | 0.557 | 0.026 | 0.229 |
| Average of Resnik Similarity | 0.558 | 0.035 | 0.219 |
| Best Match Average of Resnik Similarity | 0.528 | 0.049 | 0.222 |
| Max of SimIC | 0.555 | 0.029 | 0.228 |
| Average of SimIC | **0.573** | 0.035 | 0.235 |
| Best Match Average of SimIC | 0.554 | 0.043 | 0.219 |
| SimGIC | 0.558 | **0.050** | **0.237** |
| **Multiscale Interactome** | **0.705** | **0.091** | **0.347** |
| **Multiscale Interactome vs Best GO Baseline** | **+23.0%** | **+82.0%** | **+46.4%** |

**Supplementary Figure 5: The multiscale interactome significantly outperforms baseline methods that incorporate functional information when predicting what drugs treat a disease.** The multiscale interactome substantially outperforms all Gene Ontology (GO) baseline methods in predicting what drugs will treat a given disease. Compared to the best GO baseline method, the multiscale interactome improves AUROC by 23.0% (0.705 vs. 0.573), Average Precision by 82.0% (0.091 vs. 0.050), and Recall@50 by 46.4% (0.347 vs. 0.237). GO baselines are described in Methods. Reported values are averaged across five-fold cross-validation.

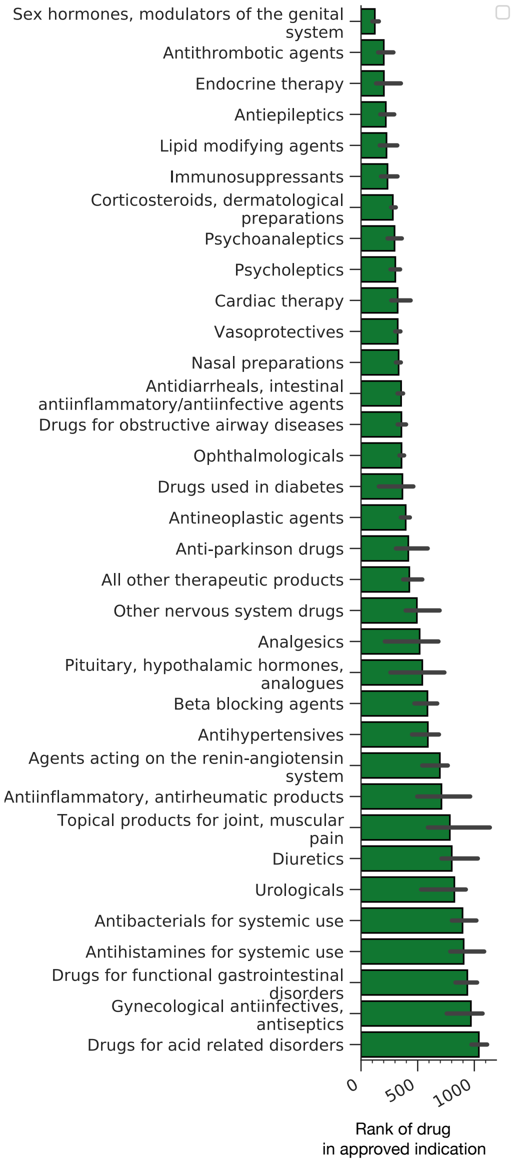


**Supplementary Figure 6: Predictive power of multiscale interactome by drug category.** The multiscale interactome predicts what drugs that treat a given disease (Figure 2a-c, Methods). For each ATC Level II drug class, the rank of the drug in its approved indication is shown (median and 95% CI).

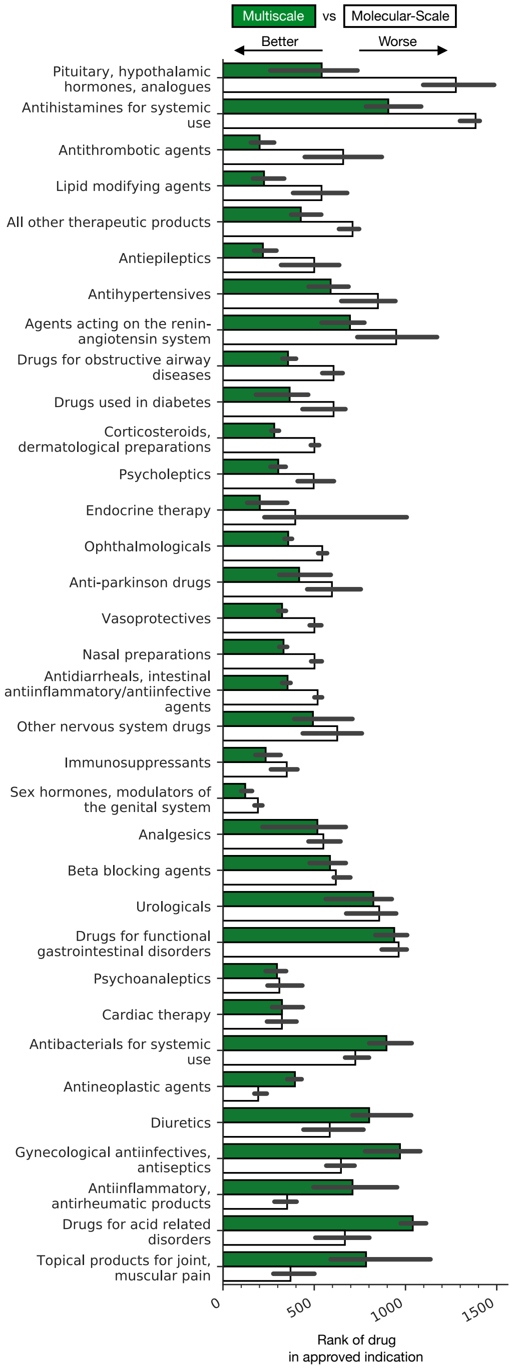


**Supplementary Figure 7: Predictive power of multiscale interactome vs molecular-scale interactome by drug category.** The multiscale interactome predicts what drug will treat a given disease more accurately than a molecular-scale interactome approach using diffusion profiles (Methods). For each ATC Level II drug class, the rank of the corresponding drugs in their approved indications is shown for both the multiscale interactome and the molecular-scale interactome (median and 95% CI). Drug classes are ordered by how much the multiscale interactome outperforms the molecular-scale interactome.

| Method | AUROC | Avg Prec | Rec@50 |
|---|---|---|---|
| Molecular-scale Interactome | **0.620** | **0.065** | **0.264** |
| Multiscale Interactome + Uberon | **0.715** | **0.091** | **0.343** |
| Multiscale Interactome + Cell Ontology | 0.710 | 0.092 | 0.329 |
| Multiscale Interactome + Uberon + Cell Ontology | 0.702 | 0.090 | 0.335 |
| Multiscale Interactome + Uberon vs. Molecular-scale interactome | **+15.3%** | **+40.0%** | **+29.9%** |

**Supplementary Figure 8: Alternative multiscale interactomes that explicitly model cells, tissues, organs, and the relationships between them also outperform molecular-scale interactomes on a drug-indication prediction task.** The best alternative multiscale interactome (multiscale interactome + Uberon) outperforms a molecular-scale interactome by 15.3% in AUROC (0.715 vs. 0.620), 40.0% in Average Precision (0.091 vs. 0.065), and 29.9% in Recall@50 (0.343 vs. 0.264). Uberon[22,23] and the Cell Ontology[24,25] are anatomical ontologies with explicit representations of cells, tissues, organs, and the relationships between them. We use human-specific subsets of both when constructing the alternative multiscale interactomes (Methods, Supplementary Note 4). Reported values are averaged across five-fold cross-validation.

| Method | AUROC | Avg Prec | Rec@50 |
|---|---|---|---|
| Multiscale interactome without GO Filtering | 0.706 | 0.092 | 0.346 |
| Multiscale interactome with GO filtering | 0.705 | 0.091 | 0.347 |

**Supplementary Figure 9: Filtering the Gene Ontology to only include biological functions associated with at least one drug target or disease protein does not substantially affect the performance of the multiscale interactome when predicting what drugs treat a disease.** Details provided in Supplementary Note 5.

| Method | AUROC | Avg Prec | Rec@50 |
| --- | --- | --- | --- |
| **Multiscale interactome without differentiating GO relation types** | 0.705 | 0.091 | 0.347 |
| **Multiscale interactome with differentiating GO relation types** | 0.709 | 0.088 | 0.335 |

**Supplementary Figure 10: Augmenting the multiscale interactome so it differentiates relation types in the Gene Ontology does not significantly improve performance on a drug-indication prediction task.** Details provided in Supplementary Note 5.

| Method | AUROC | Avg Prec | Rec@50 |
|---|---|---|---|
| Multiscale interactome without differentiating disease-protein associations types | 0.705 | 0.091 | 0.347 |
| Multiscale interactome with differentiating disease-protein associations types | 0.708 | 0.099 | 0.337 |

**Supplementary Figure 11: Augmenting the multiscale interactome so it differentiates between disease-protein associations that are based on mutation, post-translational modification, or gene expression does not substantially affect the performance of the multiscale interactome when predicting what drugs treat a disease.** Details provided in Supplementary Note 5.

| Drug-Drug Comparison Method | Spearman ρ | p-value |
|---|---|---|
| **GO Term Overlap** | 0.144 | 0.077 |
| **GSEA Term Overlap** | 0.179 | 0.028 |
| **Diffusion Profile Similarity** | **0.392** | **$5.8x10^{-7}$** |

**Supplementary Figure 12: Diffusion profiles provide a more biologically relevant metric for drug-drug similarity than alternative functional approaches.** We measure the Spearman correlation between the similarity of two drugs according to their gene expression signatures and the similarity of two drugs according to three drug-drug comparison methods: (1) intersection in GO terms associated with drug targets and disease proteins, (2) intersection in GO terms enriched among drug targets and disease proteins according to Gene Set Enrichment Analysis (GSEA)[3], and (3) similarity in diffusion profiles (Methods). The similarity of two drugs according to the similarity of their diffusion profiles provides the strongest and most significant correlation with the similarity of the same drugs according to their gene expression signatures (Spearman $\rho = 0.392$; $p = 5.8x10^{-7}$, $n = 152$; Figure 2f).

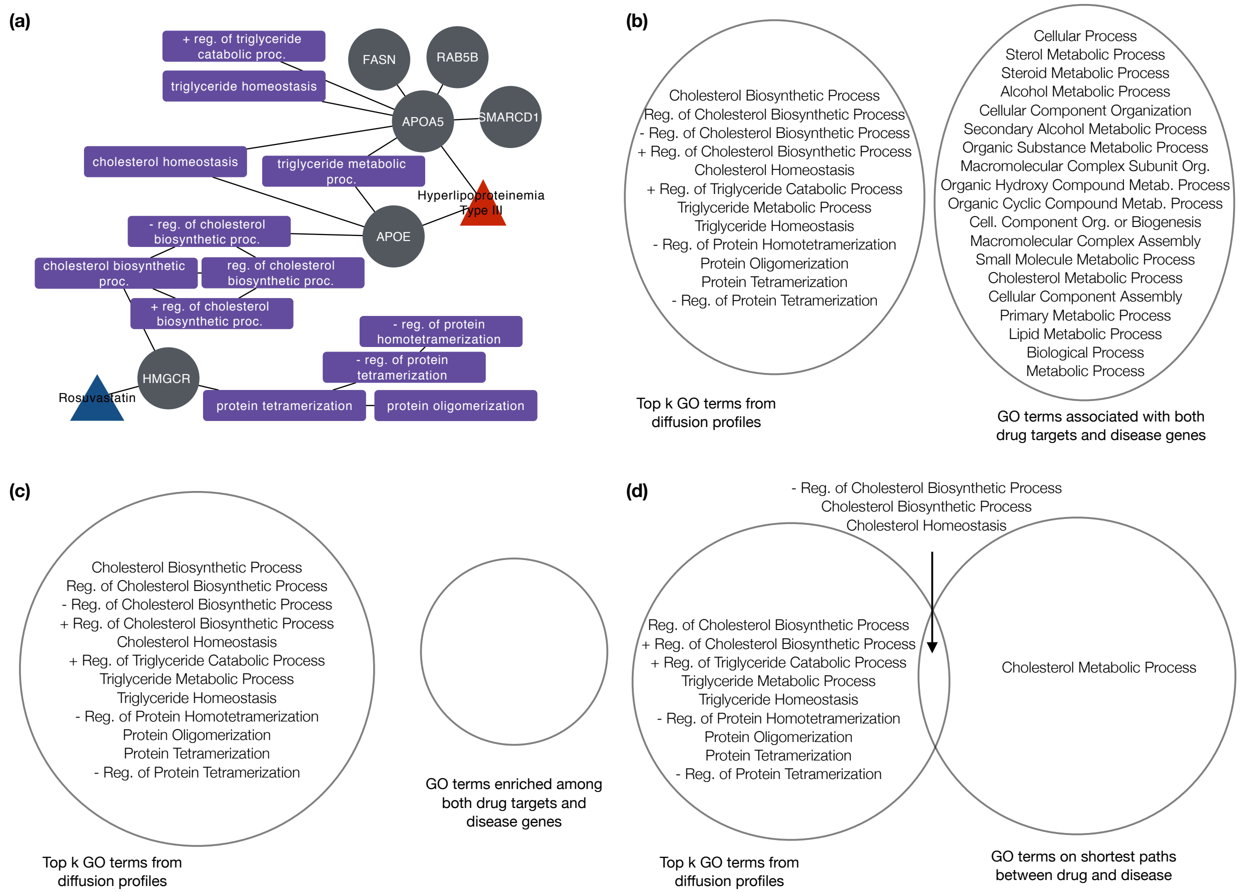


**Supplementary Figure 13: Diffusion profiles identify the biological functions used by Rosuvastatin to treat Hyperlipoproteinemia Type III more effectively than alternative approaches.** **(a)** The induced subgraph of proteins and biological functions identified via diffusion profiles as relevant to the treatment of Hyperlipoproteinemia Type III by Rosuvastatin. **(b, c, d)** Venn Diagrams compare the biological functions identified via diffusion profiles to the biological functions identified via **(b)** GO terms associated with both drug targets and disease genes, **(c)** GO terms enriched among both drug targets and disease genes according to Gene Set Enrichment Analysis[3], and **(d)** GO terms on shortest paths between drug targets and disease genes. Diffusion profiles identify the key biological functions involved in Rosuvastatin's treatment of Hyperlipoproteinemia Type III more effectively than alternative approaches (Supplementary Note 3)[11–17].

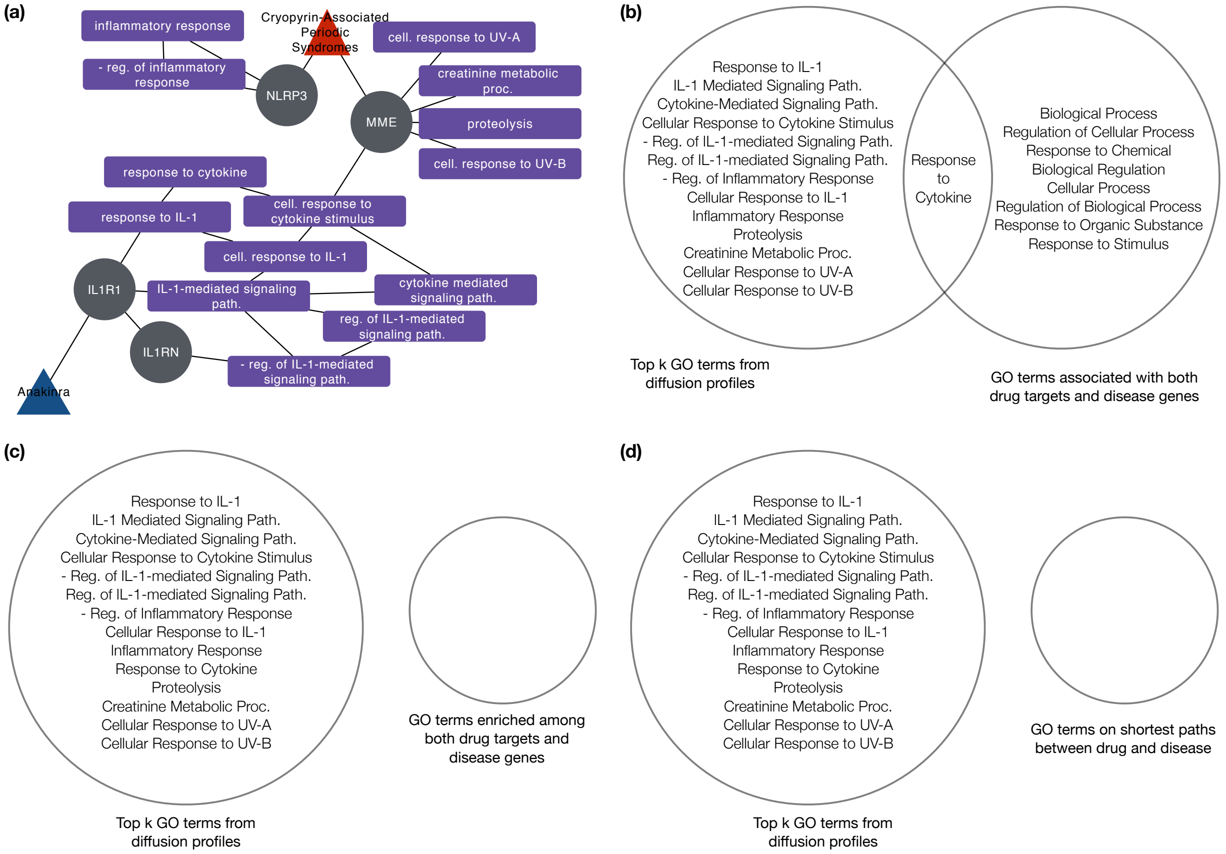


**Supplementary Figure 14: Diffusion profiles identify the biological functions used by Anakinra to treat Cryopyrin-Associated Periodic Syndromes more effectively than alternative approaches.** **(a)** The induced subgraph of proteins and biological functions identified via diffusion profiles as relevant to the treatment of Cryopyrin-Associated Periodic Syndromes by Anakinra. **(b, c, d)** Venn Diagrams compare the biological functions identified via diffusion profiles to the biological functions identified via **(b)** GO terms associated with both drug targets and disease genes, **(c)** GO terms enriched among both drug targets and disease genes according to Gene Set Enrichment Analysis[3], and **(d)** GO terms on shortest paths between drug targets and disease genes. Diffusion profiles identify the key functions involved in Anakinra's treatment of Cryopyrin Associated Periodic Syndromes – regulation of immune-mediated inflammation through the Interleukin-I beta signaling pathway[8–10]– but alternative approaches do not (Supplementary Note 3).

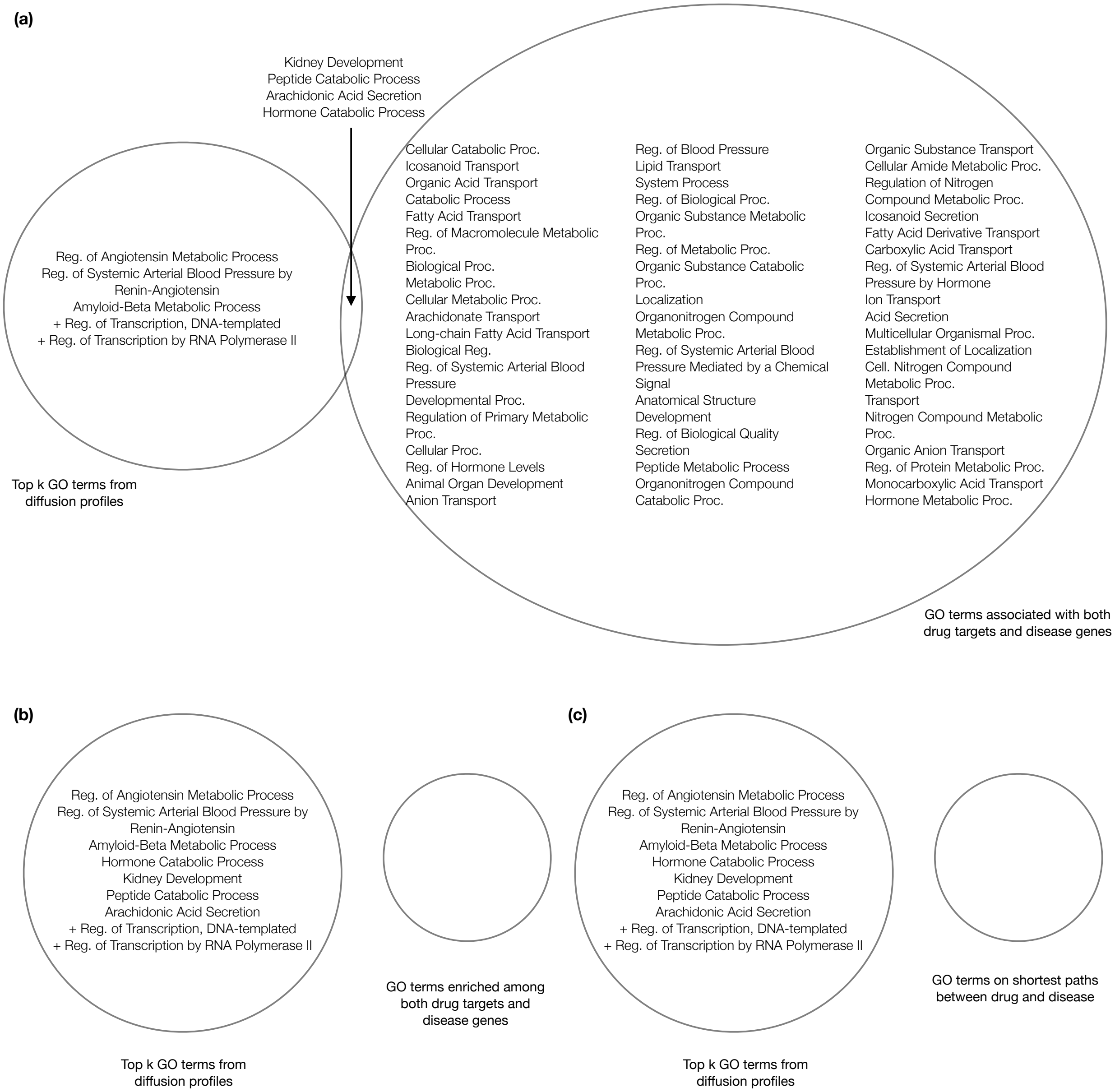


**Supplementary Figure 15: Diffusion profiles identify the biological functions used by Benazepril to treat Hypertensive Disease more effectively than alternative approaches. (a, b, c)** Venn Diagrams compare the biological functions identified as relevant to the treatment of Hypertensive Disease by Benazepril via diffusion profiles to the biological functions identified via **(a)** GO terms associated with both drug targets and disease genes, **(b)** GO terms enriched among both drug targets and disease genes according to Gene Set Enrichment Analysis[3], and **(c)** GO terms on shortest paths between drug targets and disease genes. Diffusion profiles uniquely identify the renin-angiotensin pathway as a key biological function involved in treatment, helping explain why mutations in Angiotensinogen (AGT) alter the efficacy of treatment[26–30].

| | Rosuvastatin | Hyperlipoproteinemia Type III |
|---|---|---|
| 1 | HMGCR | APOE |
| 2 | Cholesterol Biosynthetic Process | APOA5 |
| 3 | Protein Tetramerization | RAB5B |
| 4 | Protein Oligomerization | SMARCD1 |
| 5 | Negative Regulation of Cholesterol Biosynthetic Process | FASN |
| 6 | Positive Regulation of Cholesterol Biosynthetic Process | Triglyceride Homeostasis |
| 7 | Regulation of Cholesterol Biosynthetic Process | Cholesterol Homeostasis |
| 8 | Negative Regulation of Protein Homotetramerization | Positive Regulation of Triglyceride Catabolic Process |
| 9 | Negative Regulation of Protein Tetramerization | Triglyceride Metabolic Process |
| 10 | Protein Heterotetramerization | Acylglycerol Homeostasis |
| 11 | Protein Homotetramerization | Triglyceride Catabolic Process |
| 12 | Regulation of Protein Tetramerization | Positive Regulation of Lipoprotein Lipase Activity |
| 13 | FGF1 | Lipid Transport |
| 14 | Sterol Biosynthetic Process | Positive Regulation of Fatty Acid Biosynthetic Process |
| 15 | Cholesterol Metabolic Process | Positive Regulation of Lipid Catabolic Process |
| 16 | MVK | Positive Regulation of Very-Low-Density Lipoprotein Particle Remodeling |
| 17 | ERLIN2 | Osteoblast Diifferentiation |
| 18 | Secondary Alcohol Biosynthetic Process | Antigen Processing and Presentation |
| 19 | HMGCL | Chromatin Remodeling |
| 20 | PMVK | Nucleosome Disassembly |

**Supplementary Figure 16: Top 20 proteins and biological functions for the treatment of Hyperlipoproteinemia Type III by Rosuvastatin.** Proteins and biological functions are ranked according to their visitation frequency in the drug and disease diffusion profiles (Methods).

| | Anakinra | Cryopyrin Associated Periodic Syndromes |
|---|---|---|
| 1 | IL1R1 | MME |
| 2 | Interleukin-1 Mediated Signaling Pathway | NLRP3 |
| 3 | Response to Interleukin-1 | Proteolysis |
| 4 | Cellular Response to Interleukin-1 | Cellular Response to UV-B |
| 5 | Negative Regulation of Interleukin-1-Mediated Signaling Pathway | Cellular Response to UV-A |
| 6 | Regulation of Interleukin-1-Mediated Signaling Pathway | Cellular Response to Cytokine Stimulus |
| 7 | Response to Cytokine | Creatinine Metabolic Process |
| 8 | Cytokine-Mediated Signaling Pathway | Negative Regulation of Inflammatory Response |
| 9 | IL1RN | Inflammatory Response |
| 10 | IRAK1 | Positive Regulation of Interleukin-1 Beta Secretion |
| 11 | IRAK2 | Positive Regulation of NF-kappaB Transcription Factor Activity |
| 12 | ZNF675 | Negative Regulation of NF-kappaB Transcription Factor Activity |
| 13 | VRK2 | Negative Regulation of Acute Inflammatory Response |
| 14 | TRAF6 | Positive Regulation of Cysteine-Type Endopeptidase Activity Involved in Apoptotic Process |
| 15 | IRAK3 | Negative Regulation of Interleukin-1 Beta Secretion |
| 16 | MYD88 | Cellular Response to Lipopolysaccharide |
| 17 | PRKCA | Negative Regulation of NF-kappaB Import Into Nucleus |
| 18 | SRC | Cellular Response to UV |
| 19 | Cellular Response to Cytokine Stimulus | Negative Regulation of Acute Inflammatory Response to Antigenic Stimulus |
| 20 | Positive Regulation of Transcription by RNA Polymerase II | Cytoplasmic Sequestering of NF-kappaB |

**Supplementary Figure 17: Top 20 proteins and biological functions for the treatment of Cryopyrin Associated Periodic Syndromes by Anakinra.** Proteins and biological functions are ranked according to their visitation frequency in the drug and disease diffusion profiles (Methods).

| | Benazepril | Diltiazem | Hypertensive Disease |
|---|---|---|---|
| 1 | ACE | KCNA5 | Positive Regulation of Transcription by RNA Polymerase II |
| 2 | Arachidonic Acid Secretion | CACNA1C | Cilium Assembly |
| 3 | Peptide Catabolic Process | CACNA2D1 | Positive Regulation of Transcription, DNA-templated |
| 4 | Regulation of Systemic Arterial Blood Pressure by Renin-Angiotensin | CACNA1S | EGFR |
| 5 | Kidney Development | HTR3A | Negative Regulation of Transcription by RNA Polymerase II |
| 6 | Amyloid-Beta Metabolic Process | CACNG1 | ATP1A1 |
| 7 | Hormone Catabolic Process | LMNA | FN1 |
| 8 | Regulation of Angiotensin Metabolic Process | VAPA | TP53 |
| 9 | BDKRB2 | Calcium Ion Transport | LZTFL1 |
| 10 | COMT | Muscle Contraction | TGFB1 |
| 11 | AGXT | Regulation of Heart Rate by Cardiac Conduction | Negative Regulation of Transcription, DNA-templated |
| 12 | ACTB | Cardiac Muscle Cell Action Potential Involved in Contraction | APOE |
| 13 | LMNA | Negative Regulation of Cell Proliferation | CUL3 |
| 14 | EGR1 | Regulation of Calcium Ion Transport | PRKACA |
| 15 | AGR2 | Positive Regulation of Cell Aging | WNK4 |
| 16 | MYH9 | SRI | SMAD3 |
| 17 | EWSR1 | Regulation of Calcium Ion Transmembrane Transport via High Voltage-Gated Calcium Channel | TNF |
| 18 | CSNK2A2 | Atrial Cardiac Muscle Cell Action Potential | IDUA |
| 19 | AGTR2 | Potassium Ion Transmembrane Transport | CAV1 |
| 20 | CSNK2A1 | Regulation of Ventricular Cardiac Muscle Cell Membrane Repolarization | EDN1 |

**Supplementary Figure 18: Top 20 proteins and biological functions for the treatment of Hypertensive Disease by Benazepril or Diltiazem.** Proteins and biological functions are ranked according to their visitation frequency in the drug and disease diffusion profiles (Methods).
